# High Dimensional Immune Profiling Reveals CD39 as a Correlate of Tuberculosis Disease Severity

**DOI:** 10.64898/2026.07.01.735885

**Authors:** Jacqueline Watt, Dustin Sokolowski, Wenxi Xu, Ying Quan, Kyle Burrows, Jason Galutira, Blair Gordon, David G. Brooks, Jun Liu

## Abstract

Immune biomarkers of tuberculosis (TB) disease severity present a challenging area of research that remains poorly understood. New technologies are able to perform larger, unbiased studies that can unravel the complex host-pathogen dynamics occurring during a *Mycobacterium tuberculosis* infection, the causative agent of TB. In this study, we designed a high dimensional approach combining viable bacterial burden (CFU, colony forming units) with time-of-flight mass cytometry (CyTOF) analysis to profile differences in cell-type abundance and cell-type specific protein expression during states of low, intermediate and high TB disease burden. Broadly, we segregated cell-type specific immune responses into those driven by bacterial burden and/or the mycobacterial infection strain. Interrogating these immune signatures allowed us to identify ATP-catabolizing protein CD39 as a correlate of disease severity. Treatment of mice with a small molecule inhibitor of CD39 promoted effector T cell functions and CD4 T cell expansion during *Mtb* infection. Collectively, our data defines the differential lung immune environment between various mycobacterial disease severity states and uncovers a potential immune biomarker of infection and therapeutic immunomodulating target to aid in the treatment of TB.

## Introduction

The cell-mediated immunity programs that mediate tuberculosis (TB) disease severity, caused by *Mycobacterium tuberculosis* (*Mtb*) infection, remain elusive. TB has returned as the leading cause of death from an infectious agent following the COVID-19 pandemic, with an estimated 1.25 million deaths in 2023 and 10.8 million new disease cases (1). Bacille Calmette-Guérin (BCG), attenuated *Mycobacterium bovis*, is the only approved vaccine for prevention of TB. Clinical studies have confirmed an effectiveness of >80% for BCG against disseminated TB in children, however its effectiveness in adults is limited, ranging from 0-80% (2–4). Recent epidemiological models have also suggested that we may have been underestimating the portion of individuals that are able to clear or effectively control an *Mtb* infection (5). Progressing TB vaccine research has been challenging in part because there is currently no proven immunological correlate of protection for natural or vaccine-induced immunity nor biomarker for progression of disease (6,7).

The immune response to *Mtb* and the bacterial population during infection is highly dynamic and heterogeneous (8). Production of T helper cell type I (Th1) cytokines, including IFNγ and TNFα, by T cells is well documented to be critical but not sufficient for control of *Mtb* in both humans, mice and non-human primates (9). However, multiple other components including CD8 T cells, invariant natural killer T cells, γδ T cells, and CD1 restricted T cells have been shown to be involved in the cellular response to *Mtb* infection (10–12). BCG is known to induce a strong Th1 response, particularly IFNγ producing CD4 T cells, and is the only approved vaccine for TB (13). However, there are still fundamental differences between BCG and *Mtb* and immune responses they induce, which likely contributes to the ineffectiveness of BCG in adults (13,14).

We have previously described the nucleoid-associated protein Lsr2, which targets AT rich DNA and acts as a global gene regulator for ∼800 genes including many involved in *Mtb* virulence and immunogenicity, mainly in an inhibitory role (15–18). Other previous work has demonstrated Lsr2 is required for persistent infection of *Mtb* in BALB/c mice and *Mtb*’s ability to adapt to hypoxic environments (19), thus it may serve as a candidate to interrogate the immune environment under a different *Mtb* infection state.

As of now *in vitro* models are insufficient to recapitulate all the nuances of the relationship between *Mtb* and the host immune response (8). Thus, comprehensive immune profiling of an *in vivo* model of *Mtb* infection would improve the basic understanding of TB and potentially identify biomarkers for disease. Time-of flight mass cytometry (CyTOF) is a high-dimensional single-cell technology that enables the detection and quantification of many cell proteins using heavy metal polymers coupled to monoclonal antibodies (20). Recently, CyTOF has been used to study immune profile alterations in humans following TB treatment and between individuals with latent tuberculosis infection and active TB, and in mice to assess the impact of chronic viral infection on the immune response during early *Mtb* coinfection (21–24).

Here we designed a high dimensional proteomic single-cell approach to investigate the immune landscape during various mycobacterial infection states using the strains *Mtb*, *Mtb* with loss of the protein Lsr2, and BCG in mice by combining viable bacterial burden with CyTOF analysis. We defined the pulmonary cellular composition and cell states associated with disease severity. Interrogating these immune signatures revealed CD39 as a correlate of TB disease severity and uncovered a potential new therapeutic target for treatment of TB.

## Results

### Deletion of lsr2 in Mtb results in a highly attenuated strain

We constructed an *lsr2* deletion mutant of *Mtb* H37Rv (Δ*lsr2*) marked by a hygromycin-resistance cassette gene using the previously described transducing phage system (25). We then aerosol challenged C57BL/6 mice with Δ*lsr2* or *Mtb* H37Rv (Figure 1A) and measured bacterial growth in the lungs and spleen for up to 8 weeks post-infection (p.i.). Δ*lsr2* was significantly attenuated compared to *Mtb* in both the lungs and spleen (Figure 1B-C). Notedly Δ*lsr2* showed limited growth, maintaining ∼3 log_10_ lower CFU in the lungs from 2 weeks p.i. onward (Figure 1B). Lung histology analysis also identified no detectable bacilli after acid-fast staining in Δ*lsr2* samples 8 weeks p.i. (Figure 1D-I).

**Figure 1.**
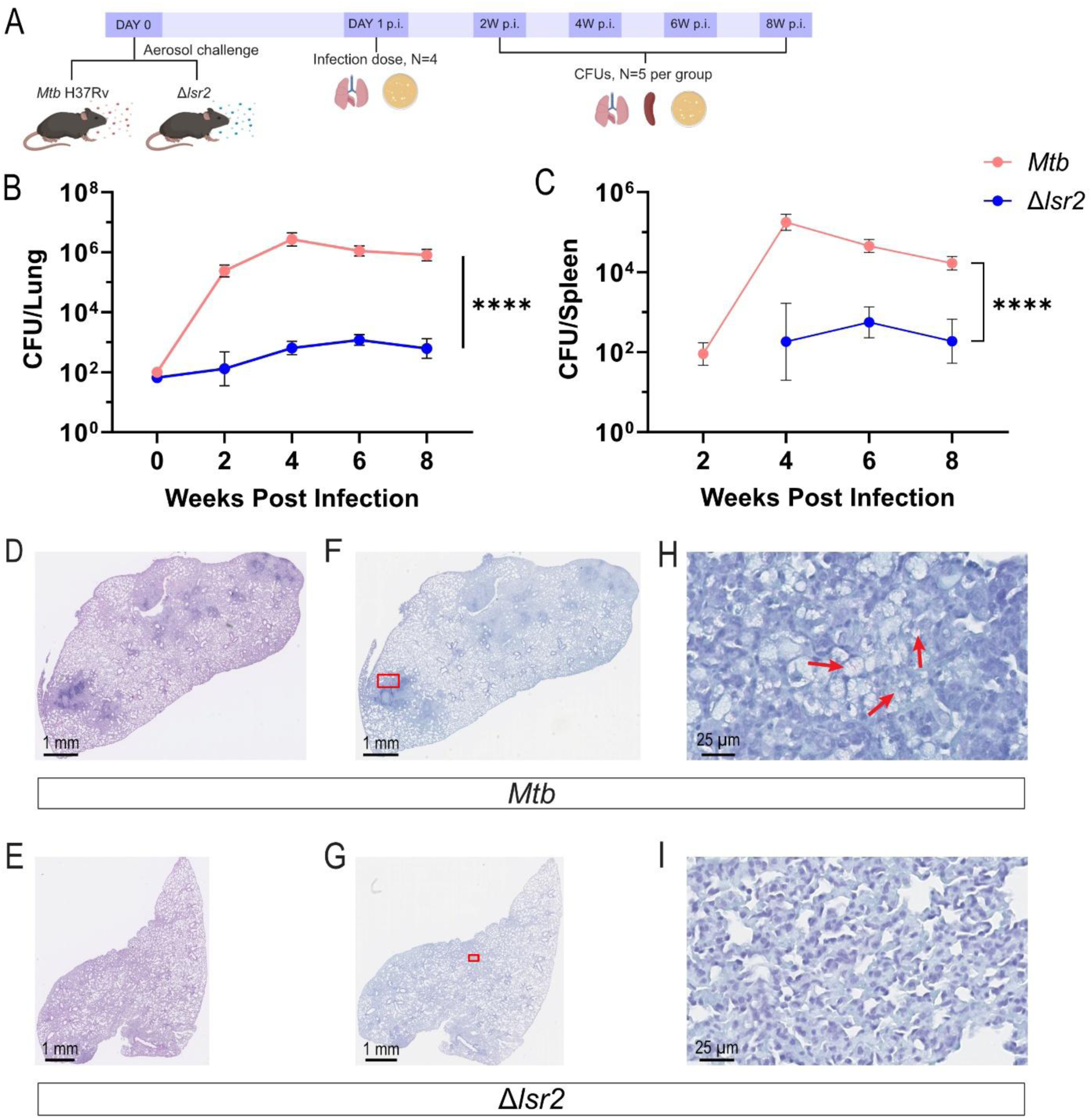
Δ*lsr2* is highly attenuated in C57BL/6 mice. (A) Schematic representation of experimental design. 6 C57BL/6 mice per group were aerosol challenged with *Mtb* H37Rv or Δ*lsr2* (∼100 CFU/lung). *Mtb* burden in the (B) lungs was determined at 2-, 4-, 6- and 8-weeks p.i. and (C) in the spleen at 2-, 4-, 6- and 8-weeks p.i. Δ*lsr2* had no detectable bacilli in the spleen at 2-weeks p.i. (Limit of Detection = 20 CFU). Data presented as mean ±SEM. Statistical significance was determined by two-way ANOVA with Sidak’s multiple comparisons test (*, p <0.05; **, p <0.01; ***, p <0.001; ****, p <0.0001; ns = not significant). Lung histological analysis was determined after 8 weeks p.i. Sections of the left lung were analyzed by (D-E) H&E staining and by (F-I) acid fast staining. Red boxes indicate area that is highlighted in neighbouring image. Red arrows indicate *Mtb* bacilli.

Δ*lsr2* is highly attenuated after aerosol challenge of mice and showed little to no sustained growth up to 8 weeks p.i. This growth profile is similar to that observed with the previously described *Mtb phoP* mutant strain SO2, which serves as the base for the live attenuated MTBVAC vaccine currently in Phase 3 clinical trials (1,14,26). Given the severe attenuation of Δ*lsr2*, we wanted to further characterize our mutant strain for expression of phthiocerol dimycocerosates (PDIMs). PDIMS are a well-studied virulence factor of *Mtb* and strains deficient in PDIM are highly attenuated in mouse models of infection (27,28). Furthermore, it is well recognized that during *in vitro* passage, *Mtb* H37Rv is highly prone to losing the ability to synthesize PDIM via spontaneous mutations (29,30). Using two-dimensional thin layer chromatography, as previously described (31), we determined that our Δ*lsr2* strain was PDIM-deficient, which likely occurred via spontaneous mutations (Figure S1). Interestingly, MTBVAC’s second attenuating mutation also disrupts the production of PDIMs; which could suggest that our Δ*lsr2* strain is a potential candidate for a live attenuated vaccine for TB.

### Profiling the mouse lung immune environment following infection with wild-type Mtb, Mtb Δlsr2, or BCG

Bacterial burden reflects TB disease severity in the mouse (32). We designed a three-way comparative model of a low, intermediate, and high bacterial burden mycobacterial infection and comprehensively profiled the cellular immune response across these various outcomes. We performed CyTOF analysis using 38 immune proteins on the adult female C57BL/6 mouse lung six weeks following aerosol infection by BCG, Δ*lsr2*, or *Mtb* H37Rv to measure the protein-level lung immune response at a single-cell resolution (Figure 2A). BCG showed little to no growth overtime, with a bacterial burden similar to the inoculation dose of ∼100 CFU (Figure 2B); thus, it represents low TB disease severity. We took advantage of the severe attenuation of Δ*lsr2*, with a moderate bacterial burden ∼1 log_10_ higher than BCG (Figure 2B), to represent intermediate TB disease severity. The parental strain *Mtb* H37Rv had a bacterial burden of ∼3 log_10_ higher than Δ*lsr2* (Figure 2B) and represented a high TB disease severity state. Using these three groups we aimed to dissect immune features related to virulence and disease progression. Immune proteins interrogated include those associated with cell-type, activation, and immune checkpoints.

**Figure 2.**
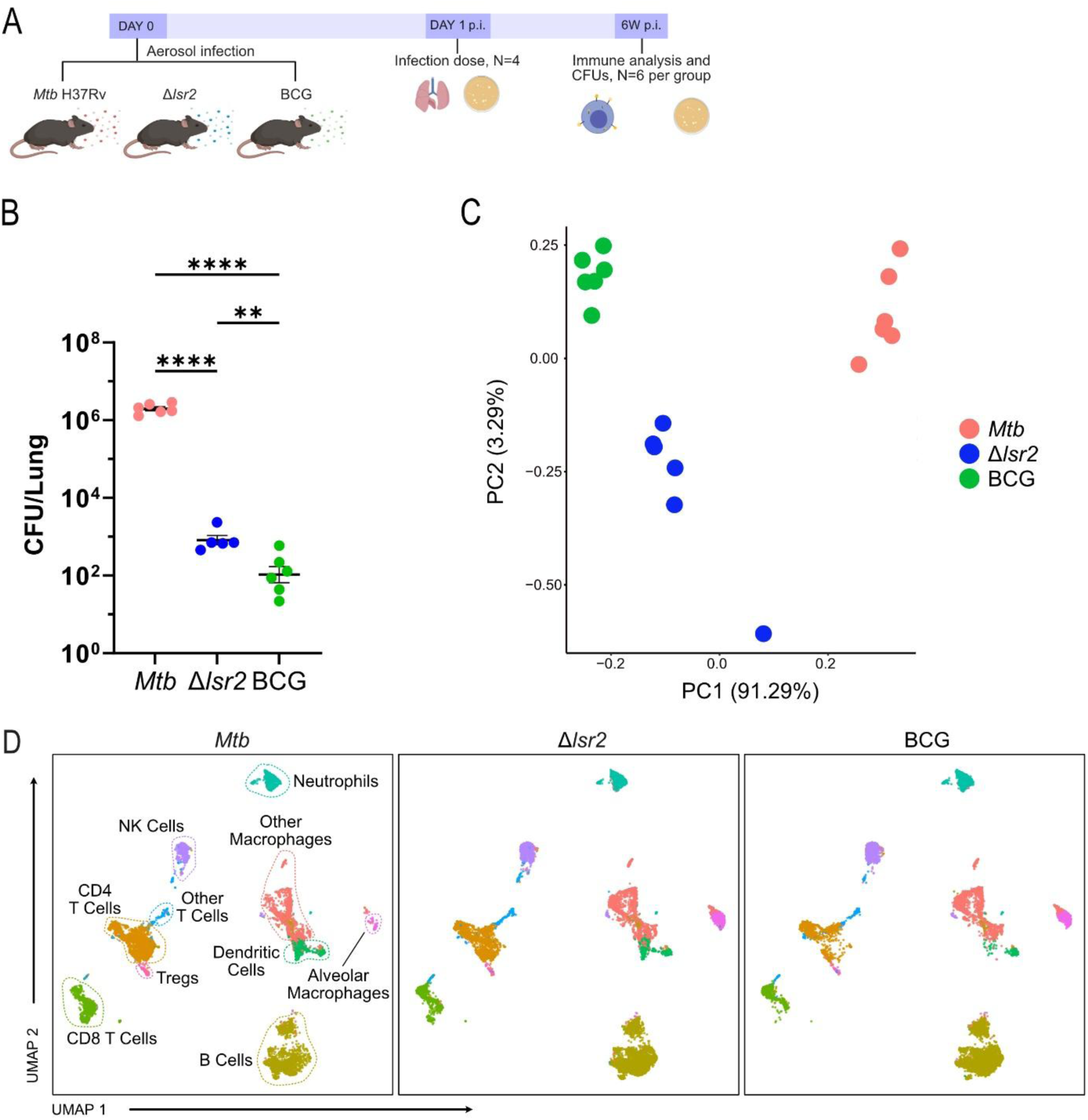
Global summary of the lung immune system 6 weeks after *Mtb*, Δ*lsr2*, or BCG infection. (A) Schematic of experimental design. 6 C57BL/6 mice per group were aerosol challenged with *Mtb* H37Rv, Δ*lsr2*, or BCG (∼100 CFU/lung). *Mtb* burden in the (B) lungs was determined at 6 weeks p.i. Lung single cell suspensions from non-perfused lungs were analyzed with CyTOF. (C) Scatterplot visualizing the first two principal components of pseudobulked cell-type specific protein expression. Colours represent condition. (D) Uniform manifold approximation mapping (UMAP) of cell-type specific protein expression. Panels are split by biological condition. Cells are labelled by cell-type identified by manually annotating metaclusters based on their protein expression profiles.

After quality control and filtering, we collected 1,696,846 total cells (mean 94,269 cells per sample) whose cell-type identity and cell-state could be profiled. Pseudobulked PCA showed that the three groups were linearly separable across PC1 which accounts for 91.29% of the variation in this dataset, with BCG samples clustering more closely to Δ*lsr2* samples than to *Mtb* samples (Figure 2C). A total of 10 clusters were identified, comprising the major immune cell subsets (Figure 2D).

We profiled differences in cell-type abundance and cell-type specific protein expression between the three groups and classified them into one of three states: (1) CFU: differential based on bacterial burden, low TB disease severity vs. intermediate TB disease severity vs. high TB disease severity; (2) ATTN: differential based on attenuated strains, BCG and Δ*lsr2* vs. *Mtb* H37Rv; (3) MTB: differential based on genetic similarity, *Mtb* H37Rv and Δ*lsr2* vs. BCG.

Eight cell-types were differentially abundant between *Mtb* and the other lower bacterial burden groups (B cells, other macrophages, alveolar macrophages, CD4 T cells, regulatory T cells [Tregs], other T cells, CD8 T cells, and NK cells), making cell-type composition broadly conditional on attenuated bacterial strains (ATTN; Figure 3A). Neutrophil abundance was stable across groups. B cell proportions and alveolar macrophage proportions were significantly reduced in the *Mtb* group (Figure 3B). *Mtb* infection also exhibited the highest proportion of CD4 and CD8 T cells in the lung (Figure 3B). These cell-type dynamics represent a well-documented pan-immune reprogramming of *Mtb* infection (33). Interestingly, dendritic cells displayed a CFU pattern, incrementally decreasing in cell-type proportion from low TB disease severity to high TB disease severity. Dendritic cells have previously been considered an important target for *Mtb* vaccinations as they facilitate the activation of BCG-treated T-cells *in vivo* (34).

**Figure 3.**
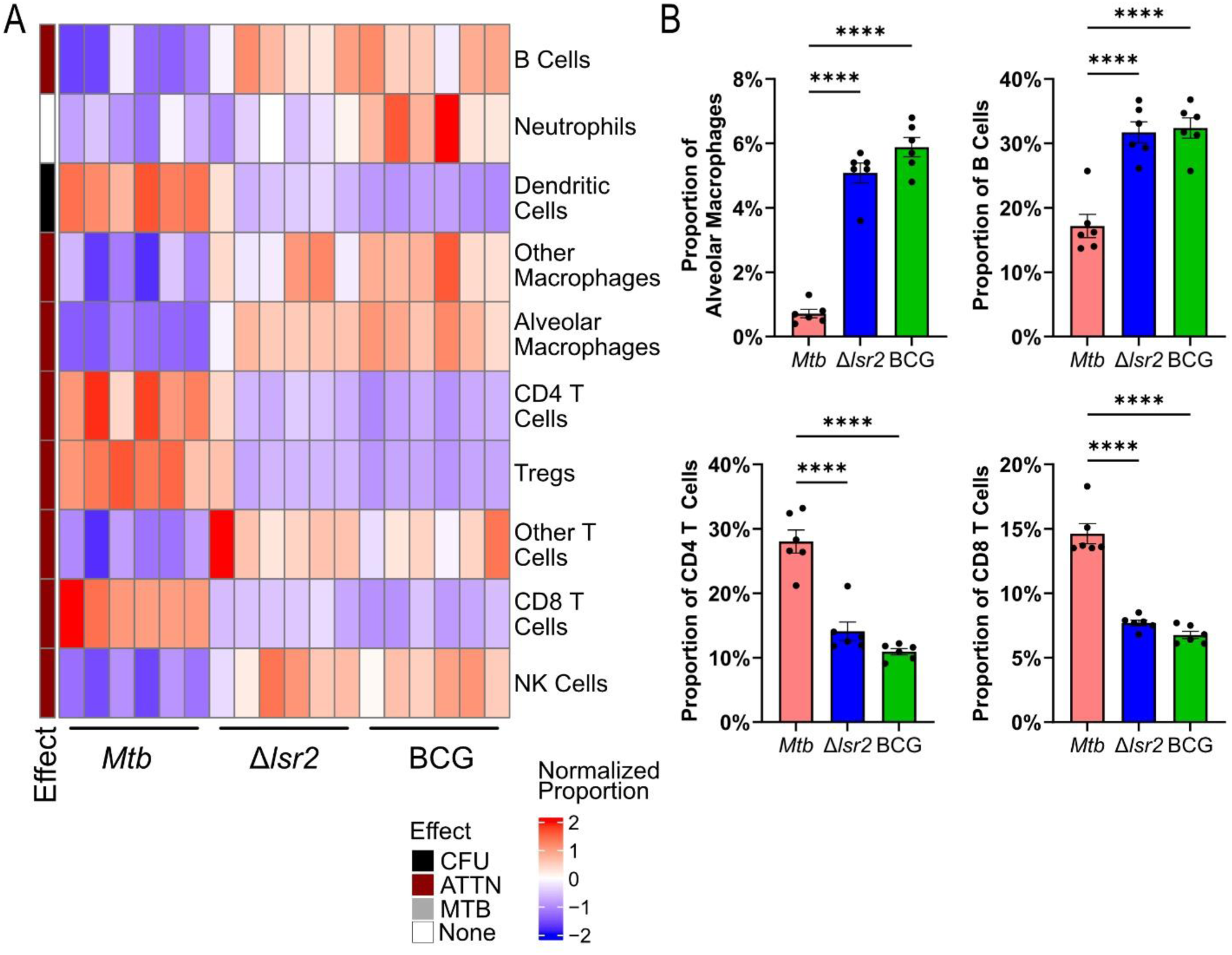
Cell-type abundance observed across TB disease severity. (A) Heatmap represents differences in cell-type abundance between each group. Rows are different cell-types, columns are different samples, and heat represents row-normalized cell-type abundance. Rows are annotated by their state (i.e. CFU, ATTN, or MTB) for cell-type differences with an FDR-adjusted p-value <0.05. None = not significant. (B) Percentage of alveolar macrophages, B cells, CD4 and CD8 T cells per sample. Numerical representation of select cell-type proportions displayed in heatmap (A) above. Differences in cell-type proportions were measured using a one-way ANOVA and Tukey’s multiple comparisons test (*, p <0.05; **, p <0.01; ***, p <0.001; ****, p <0.0001; ns = not significant).

### CD39 is overexpressed during high bacterial burden Mtb infections and is prominent on T cells

Interestingly, we observed that CD39 was a top-scoring protein for PCA-based non-redundancy scores during our CyTOF analysis (Figure S2). CD39 is an enzymatic cell surface protein that hydrolyses extracellular ATP (eATP) into adenosine (35). eATP is released by stressed or dying cells and provides inflammatory signals that are critical for effective immune responses (35). By contrast, adenosine is a potent suppressor of immune activity (36).

Upon further interrogation, we observed that all cell-types analyzed displayed high levels of CD39 expression in *Mtb* infected lung samples where bacterial burden was high (Figure 4A). Moreover, CD39 displayed an increasing CFU effect in dendritic cells, Tregs, other T cells, CD8 T cells and CD4 T Cells (Figure 5A).

**Figure 4.**
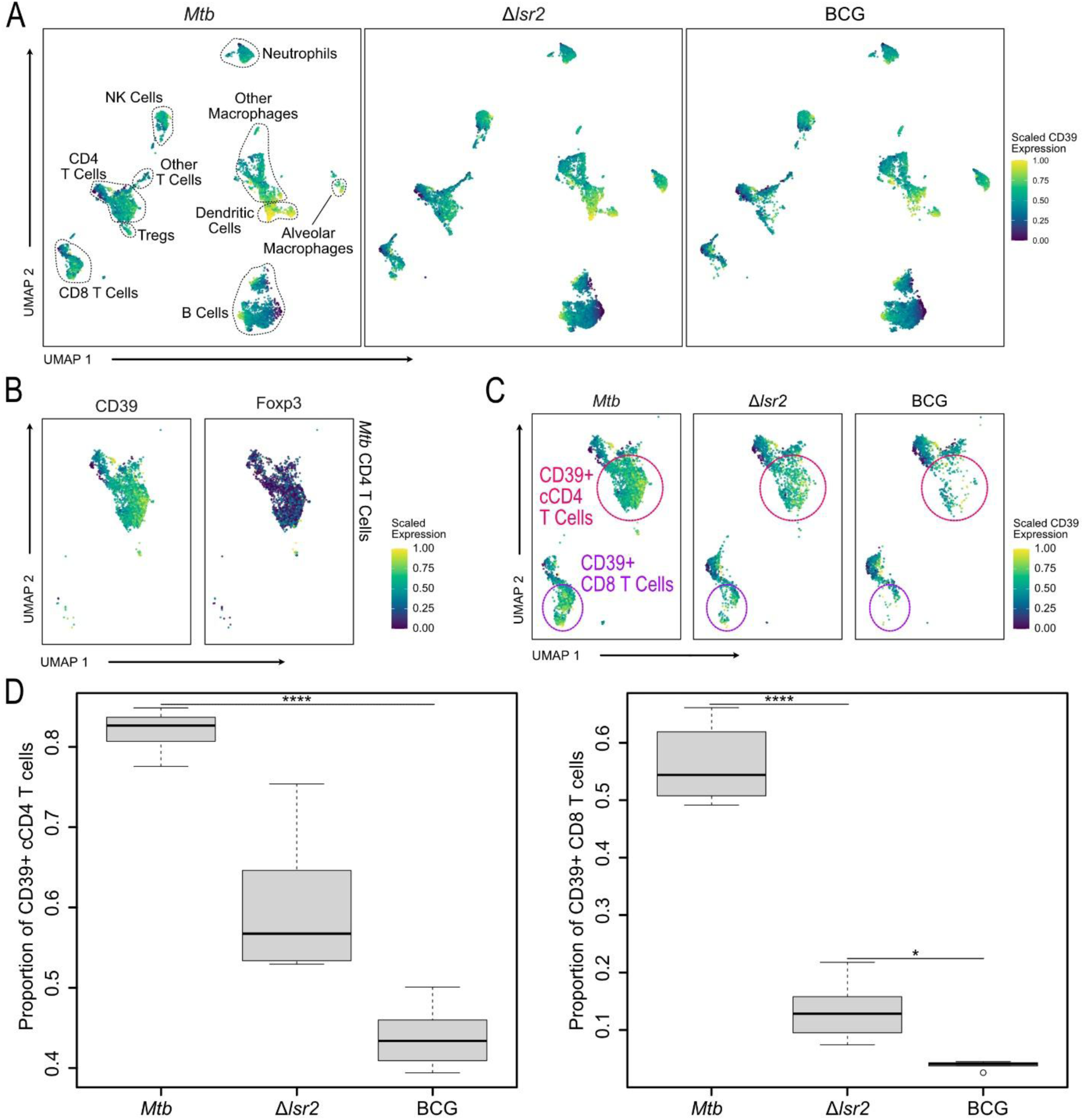
A high proportion of CD4 and CD8 T cells express CD39 after an *Mtb* infection. (A) Uniform manifold approximation mapping (UMAP) of CD39 protein expression in all cell types. (B) UMAP of protein expression in conventional CD4 (Foxp3-; cCD4) T cells in *Mtb* lung samples. Columns represent cell-type protein expression for CD39, and Foxp3. (C) UMAP of CD39 protein expression in CD4 and CD8 T cells. CD39+ cCD4 T cells are indicated by the pink circles and CD39+ CD8 T cells are indicated by the purple circles. (D) Bar plots comparing the ratio of CD39+ cCD4 T cells vs CD39- cCD4 T cells and CD39+ CD8 T cells vs CD39- CD8 T cells within each group. Differences in cell-type CD39 positivity were measured using a one- way ANOVA and Tukey’s HSD test, with each mouse as a biological replicate (*, p <0.05; **, p <0.01; ***, p <0.001; ****, p <0.0001; ns = not significant).

**Figure 5.**
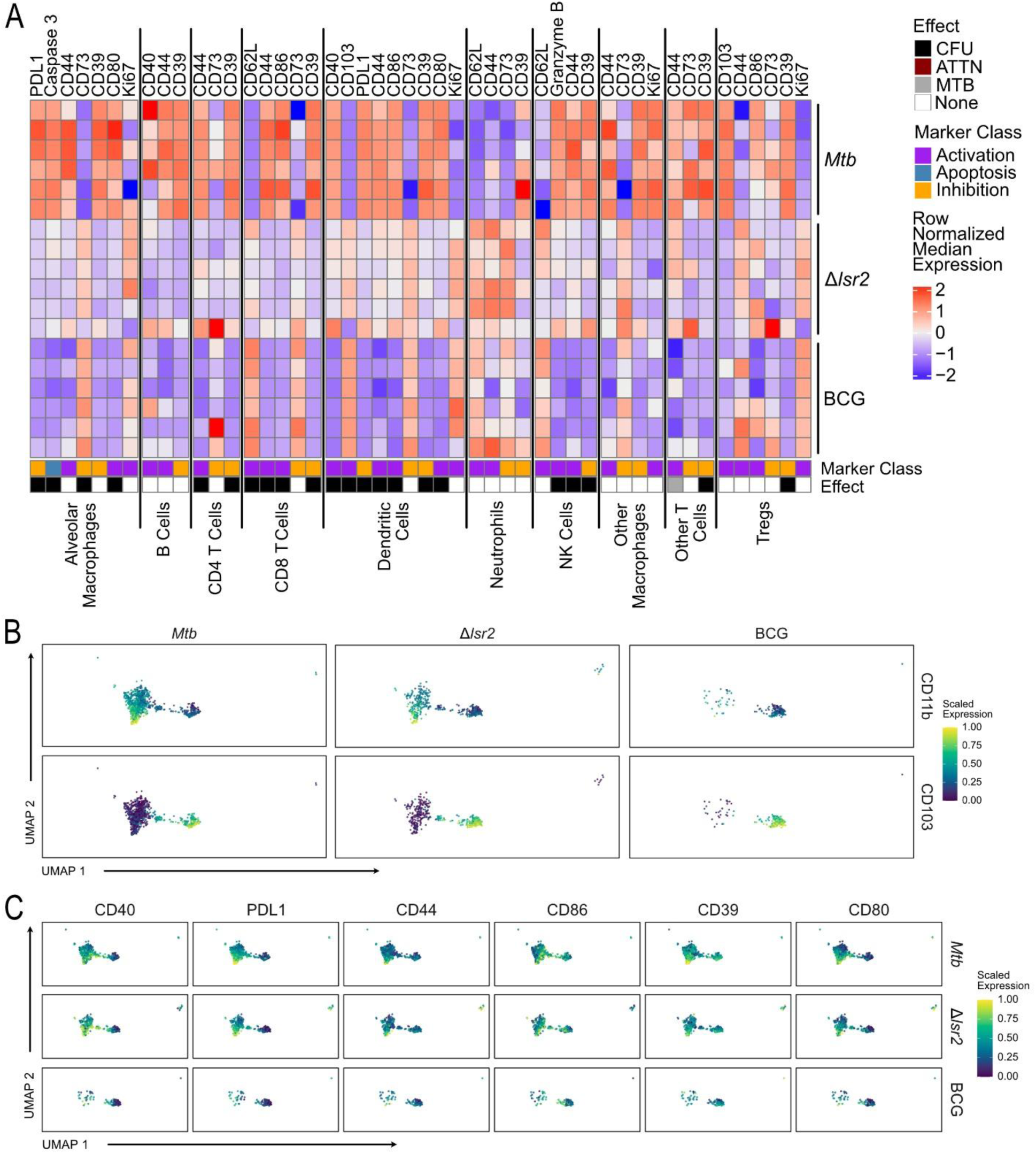
Cell-type marker expression dynamics observed across TB disease severity. (A) Heatmap represents differences in protein expression associated with cell-state between each treatment in all immune cells. Columns represent cell-type proteins and rows represent samples. Cells are populated by row-normalized protein expression. Columns are annotated by cell-type, state (i.e. CFU, ATTN, or MTB), and protein class (i.e., activation, exhaustion etc.). State is shown for cell-type protein expression differences with an FDR-adjusted p-value <0.05. None = not significant. (B-C) Uniform manifold approximation mapping (UMAP) describing dendritic cell subtypes and differentially expressed proteins. (B) Subtypes are defined by CD11b and CD103 cell-type protein expression. Columns are annotated by infection group. (C) Columns represent cell-type protein expression for CD40, PDL1, CD44, CD86, CD39, and CD80. Rows are annotated by infection group.

Given the higher abundance of CD4 and CD8 T cells in *Mtb* samples previously observed (Figure 3B), we wanted to further investigate the relationship between disease severity and CD39 expression on T cells. Interestingly, we found that CD39 is expressed on Foxp3- CD4 T cells (Figure 4B), indicating that the high expression of CD39 is being driven by conventional CD4 (cCD4) T cells not Foxp3+ Tregs (37). Therefore, we annotated cCD4 T cell subclusters as CD39+ cCD4 T cells or CD39-cCD4 T cells (Figure 4C). We also annotated CD39+ and CD39- CD8 T cells for further analyses (Figure 4C). A significantly larger population of cCD4 T cells and CD8 T cells expressed CD39 in *Mtb* infected samples but not in BCG and Δ*lsr2* infected samples (Figure 4D). To our knowledge, this is the first time such CD39+ T cell populations have been described in the mouse model of *Mtb* infection.

### Immune cell activation-exhaustion profiles in Δlsr2 infected mice resemble those of high burden Mtb infection

We next sought to investigate overall cell-type specific protein expression between groups. We focused on cell proteins associated with activation (i.e., CD103, CD44, CD86, Ki67, CD62L, GranzymeB, CD80, CD40), exhaustion (i.e., PDL1, CD39, CD73), and apoptosis (caspase 3) between conditions at a cell-type resolution.

We observed a highly dynamic T cell environment, reflecting T cells as critical responders to *Mtb* infection. We found an increasing CFU effect for CD8 T cells with CD44, and CD86 and a decreasing CFU effect of CD62L (Figure 5A).

Alveolar macrophages exhibited an increasing CFU effect of PDL1, caspase 3, and CD80 and a decreasing CFU effect of CD73, which is paired with a depletion of non-exhausted alveolar macrophages in the *Mtb* group (Figure 5A). Here, Δ*lsr2* infected mice reflect the exhaustion and apoptosis of macrophages found in *Mtb* infected mice, but to a lesser degree.

The co-expression of many proteins associated with activation and exhaustion was also found in dendritic cells. We found an increasing CFU effect in CD40, PDL1, CD44, CD86 and CD80, and a decreasing CFU effect in CD103 (Figure 5A). We found two populations of dendritic cells, defined by CD11b+ and CD103+, respectively (Figure 5B). The differential proteins mapped to the CD11b+ dendritic cell population (Figure 5C). Δ*lsr2* and *Mtb* dendritic cell populations display a diverse co-expression mixture of proteins associated with activation (e.g., CD40+) and exhaustion (CD39+, PDL1+) that are lacking in BCG infected mice. These findings suggest that Δ*lsr2* stimulates dendritic cells better than BCG and more closely reflects the dendritic cell response found during *Mtb* infection.

### Analysis of lung-infiltrating cells improves detection of immune profile differences between lower severity disease conditions

We first chose to perform our analysis on non-perfused lungs containing both circulating and non-circulating cells, however, by focusing on perfused lung samples we may obtain a more lung tissue specific immune profile to mycobacterial infection.

Accordingly, we repeated the CyTOF experiment using a 28 immune protein panel on PBS-perfused adult female C57BL/6 mouse lungs six weeks following aerosol infection by BCG, Δ*lsr2* or *Mtb* H37Rv. In this experiment, after quality control and filtering, we collected 856,867 total cells (mean 47,603) cells per sample. A total of 10 immune cell subsets were identified (Figure S3A). Like the non-perfused experiment, pseudobulked PCA showed that inoculated groups were linearly separable across PC1 which accounts for 79.39% of the variation in this dataset, however unlike in the previous dataset, all three groups were spaced equidistantly (Figure S3B).

Notedly, we again found an increasing CFU effect of CD39 expression in CD4 T cells, supporting our findings from the first dataset of non-perfused lung samples (Figure S4B-C). We also saw an increasing ATTN effect of CD44 expression in CD4 T cells and an increasing CFU effect of CD44 expression in CD8 T cells, suggesting a similar degree of T cell activation in Δ*lsr2* and *Mtb* mice.

On top of recapitulating many of the results from our previous dataset, by focusing on PBS-perfused lung tissue immune cells we were able to better detect differences between the lower disease severity groups (Δ*lsr2* and BCG) and further assess functional diversity in the immune responses. We found a decreasing CFU effect of cell-type composition in macrophages and B cells, and an increasing CFU effect in CD4 T cells (Figure S3C). Unlike in the previous non-perfused dataset, we detected an increasing MTB effect in dendritic cells, where Δ*lsr2* and *Mtb* had a higher abundance of dendritic cells than BCG. Lastly, we found an ATTN effect with cell-type composition in other macrophages (decreasing), other T cells (decreasing), and CD8 T cells (increasing). We did not see a difference in monocyte, neutrophil, or NK cell composition (Figure S3C). With cell-type-specific protein expression in the PBS-perfused lung, we observed a CFU effect for alveolar macrophage expression of CD44 and CD73 (decreasing), as well as CD39 (increasing; Figure S4A). Dendritic cells had a similar co-enrichment of proteins associated with activation and exhaustion to the non-perfused lung samples. However, unlike in the previous dataset, we were able to detect a more distinct increasing CFU effect in PBS-perfused lungs of CD44 and CD73 expression, as well as a decreasing CFU effect of CD39 expression (Figure S4A and D).

### CD39 expression is highly associated with Mtb infected cells

CD39 is encoded by gene *entpd1* (ectonucleoside triphosphate diphosphohydrolase-1) (35). To spatially localize expression of *entpd1* mRNA and validate our previous CD39 protein expression observations, we performed RNAscope hybridization with automated quantification (HALO, Indica Labs) on PBS-perfused lung samples collected from the previous CyTOF experiment at the 6-week p.i. time point for *Mtb*, Δ*lsr2*, and BCG infected mice (Figure 6B). Cells bordering potential vascular structures were removed from the analysis to reduce bias from lung endothelial cells, which constitutively express CD39 (35). Colocalizing e*ntpd1* with c*d3e* (T cells) and IS6110 (*Mtb*/BCG) (Figure 6A), we observed a progressive increase in the number of e*ntpd1*+ T cells, normalized per 10^5^ cellular nuclei, from BCG to Δ*lsr2* and Δ*lsr2* to *Mtb* (Figure 6C). The total number of *entpd1*+ cells was also higher in *Mtb* and Δ*lsr2* lung samples compared to BCG (Figure 6C-D). Strikingly, *entpd1*+ cells were highly associated with *Mtb* infected cells, regardless of the infection strain (Figure 6C). The high proportion of *entpd1*+ *Mtb* infected cells further supports a direct link between *Mtb* infection and CD39 expression.

**Figure 6.**
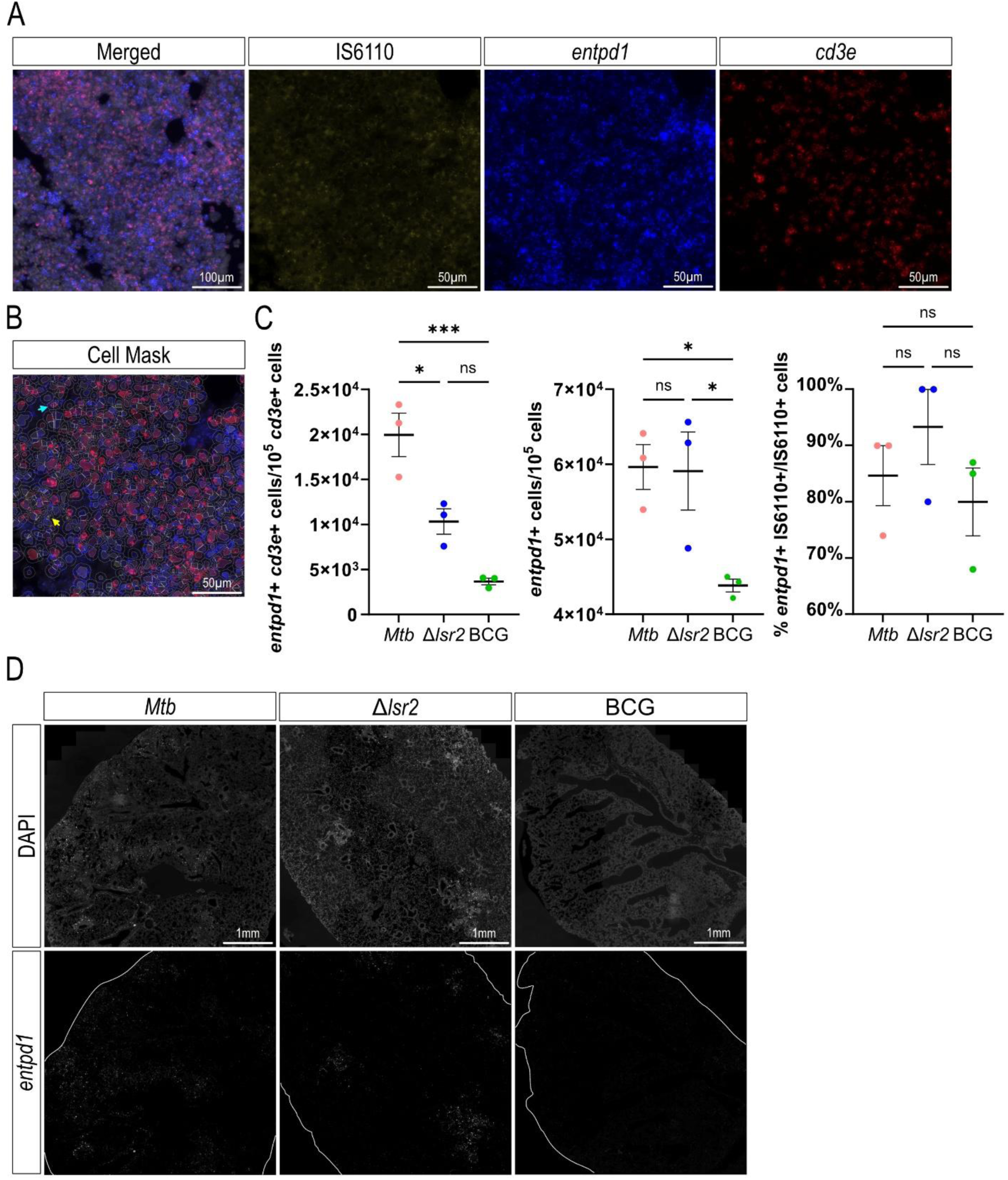
Spatial localization of CD39 expression in *Mtb*, Δ*lsr2* and BCG infected mouse lungs. (A) Representative microscopic images of RNAscope of *entpd1* (CD39, blue) with T cells (*cd3e*, red) and *Mtb*/BCG (IS6110, yellow). Images on the left show RNAscope in the lung with nuclear counterstain (DAPI, grey). Magnifications are shown on the right with the indicated staining. Scale bars represent 100 μm in the merged image and 50 μm in the magnifications. (B) Representative cell mask image output from HALO FISH module (Indica Labs). Teal and yellow arrows indicate an *entpd1*+/*cd3e*+ or *entpd1*+/IS610+ cell, respectively. Scale bar represents 50 μm. (C) Lungs from 3 randomly selected mice per group were used for automated image analysis (FISH module v3.2.3, HALO, Indica Labs). The number of *entpd1*+ *cd3e*+ per 10^5^ *cd3e*+ cells, *entpd1*+ cells per 10^5^ cells and the percentage of cells expressing *entpd1* that also express IS6110 per all cells expressing IS6110 were quantified. Data presented as mean ±SEM. Statistical significance was determined by one-way ANOVA with Tukey’s multiple comparisons test (*, p <0.05; **, p <0.01; ***, p <0.001; ****, p <0.0001; ns = not significant). (D) Representative image data for RNAscope of *entpd1* in an *Mtb*, Δ*lsr2* and BCG infected mouse lung. The top row shows the indicated lung sample with nuclear counterstain (DAPI, grey). The bottom row shows *entpd1*+ cells with the lung outlined in white. Scale bar represents 1 mm.

### Treatment with a CD39 inhibitor increases IFNγ and TNF during Mtb infection

Previous studies have demonstrated that increased CD39 expression is associated with T cell exhaustion (38–40). We next sought to investigate the role of CD39 during *Mtb* infection and the potential therapeutic impact of inhibiting CD39. Polyoxymetalate-1 (POM-1) is a potent chemical inhibitor of CD39 and has shown promising anti-tumour activity through modulation of the tumour immune environment (41,42). C57BL/6 mice were aerosol challenged with *Mtb* H37Rv for 1-, 2-, or 3-weeks and then treated with POM-1 via intraperitoneal injection for 14 days at 5 mg/kg or PBS as a control (Figure S5A), following a previously published protocol (41). It is well established that T cell infiltration to the lungs is delayed by approximately 2 weeks after *Mtb* infection (43–45). Therefore, POM-1 treatment timing was selected to encompass various periods before, during, and after T cells begin infiltrating the lungs p.i. Bacterial burden was examined in the lungs at 3 days, 2 weeks and 4-weeks post treatment (p.t.) with POM-1.

POM-1 had no major impact on the bacterial burden in the lungs or spleen under these experimental conditions (Figure S5B). These results were unchanged by the duration between *Mtb* infection and initiation of POM-1 treatment (Figure S5C-D).

T cell response and Th1-type cytokine response was assessed in mice treated with POM-1 3-weeks after *Mtb* infection. 3 days post treatment, POM-1 treated mice had increased production of TNF in the lungs in all CD45+ cells, CD4 and CD8 T cells compared to PBS control mice (Figure 7A). TNF production remained increased up to 4 weeks p.t. in CD45+ cells. We also observed a significant increase in IFNγ as well as an increasing trend of IL-2, IL-10 and IL1b expression in CD45+ cells in the lungs up to 4 weeks p.t. (Figure 7C). A greater proportion of IFNγ+ CD4 T cells was observed at 2 weeks p.t. (Figure 7B) and was maintained up to 4 weeks p.t. (Figure 7C). This finding was paired with a greater proportion of IFNγ+ CD8 T cells compared to PBS control mice (Figure 7C) and an overall increase in the proportion of CD4 T cells 4 weeks p.t. (Figure 7D). These data suggest that inhibiting CD39 may promote effector T cell functions and CD4 T cell expansion during *Mtb* infection.

**Figure 7.**
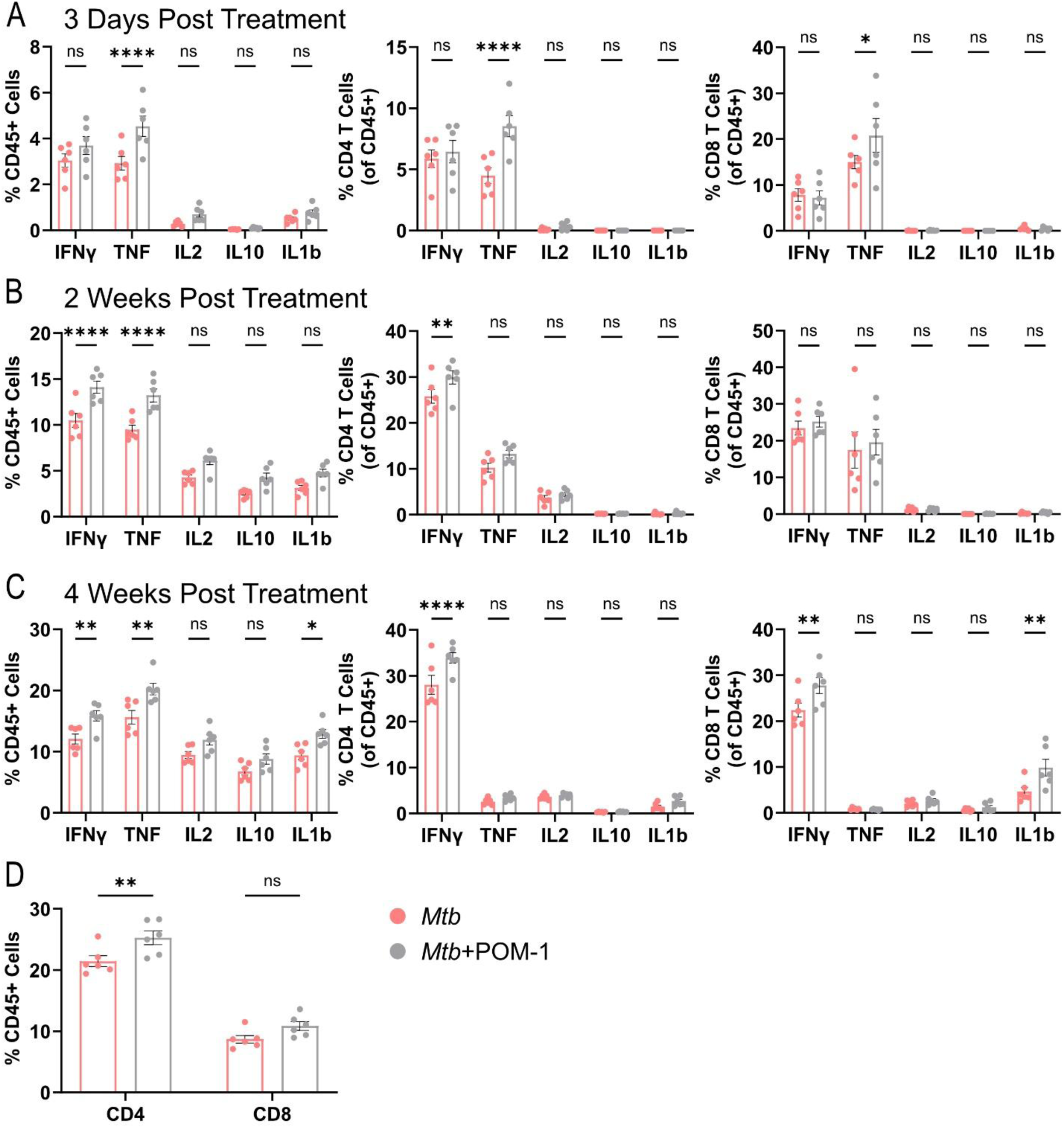
POM-1 treatment modulates the Th-1 type immune response to *Mtb* infection. C57BL/6 mice were aerosol challenged with *Mtb* H37Rv (∼100 CFU/lung) and after 3 weeks were treated with either 5 mg/kg of POM-1 or PBS as a control. Lung single cell suspensions were treated for 5 hours with eBioscience Cell Stimulation Cocktail (plus protein transport inhibitors) to determine differences in the proportion of Th-1 cytokine production at (A) 3 days post treatment (p.t.), (B) 2 weeks p.t. and (C) 4 weeks p.t. in CD45+ immune cells, CD4 T cells, and CD8 T cells. (D) Overall proportion of T cell populations at 4 weeks p.t. Data presented as mean ±SEM. Statistical significance was determined by two-way ANOVA with Sidak’s multiple comparisons test (*, p <0.05; **, p <0.01; ***, p <0.001; ****, p <0.0001; ns = not significant).

## Discussion

Discovering human TB disease relevant biomarkers in the mouse model of *Mtb* infection, the most widely used animal model for TB, would be a very useful tool to more effectively identify potential prevention and control strategies for TB. Here, we developed a mouse model comparing the immune environment during a low, intermediate and high TB disease burden representation using BCG, an attenuated strain of *Mtb* with the loss of the protein Lsr2, and the parental strain *Mtb* H37Rv, respectively. This work provides proof of concept for an approach to discover potential disease relevant biomarkers of *Mtb* infection and we also propose Δ*lsr2* as a candidate live attenuated vaccine for TB.

We developed a comparative immune analysis model that recapitulated many known immune cell dynamics observed with *Mtb* infection, in both the non-perfused and PBS-perfused lung. The stepwise decrease in macrophage composition with increasing CFU could relate to the well-studied hijacking of macrophages by *Mtb* to replicate and influence their cell death via necrosis (46). We also observed a decrease in B cell abundance as CFU increased; this is consistent with B cell depletion previously reported in human patients infected with *Mtb* (47).

Our data revealed a highly heterogenous dendritic cell response particularly in CD11b+ dendritic cells in Δ*lsr2* and *Mtb* infected mice. CD11b+ dendritic cells have been shown to play a key role in mediating Th1 cell responses against *Mtb* infection (48). Interestingly, Lai *et al.* also established that CD103+ dendritic cells counter regulated Th1 activation by inhibiting CD11b+ dendritic cell mediated Th1 cell priming via IL-10.

We also observed that BCG infected mice had the highest median expression of CD103 in dendritic cells compared to Δ*lsr2* and *Mtb*. This could indicate the dendritic cell response to BCG is skewed towards being immune-suppressive and may contribute to its inability to effectively protect against TB. Moreover, BCG exposure has previously been reported to enhance the production of IL-10 by dendritic cells (49). Taken together, our work recapitulated many of the immune responses detailed in previous TB studies but within a single controlled experimental context.

Increased abundance of CD4 T cells following *Mtb* infection is well established and has made CD4 T cells a priority target for *Mtb* vaccines, however, the specific relationship between CD4 T cells, *Mtb* and protection remains unclear (50). We identified a high expression of CD39 on all immune cells interrogated, and particularly higher expression on both CD4 and CD8 T cells following *Mtb* infection compared to Δ*lsr2* and BCG. CD39 is the rate limiting enzymatic cell surface protein that hydrolyses extracellular ATP (eATP) into AMP, which is then converted into adenosine in the purinergic signaling pathway (35). CD39 has been identified as a marker of T cell exhaustion for conventional CD4 T cells and CD8 T cells in chronic viral infections and cancer (38–40). Exposure to persistent antigen and/or inflammatory signals causes the deterioration of T cell function, a state called exhaustion (51,52). T cell exhaustion is associated with inefficient control of persisting viral infections and tumours. We hypothesize that during a high burden *Mtb* infection the high levels of cell death and disease pathology leads to the upregulation of CD39 on T cells resulting in their dysfunction and inability to control the infection. Alternatively, CD39 may be a readout of exhaustion and dysfunction in these T cells, allowing CD39 to serve as a biomarker of disease severity. Recently, patients with TB were reported to display significantly higher levels of CD39 expression on Th1 cells compared to healthy controls, further supporting our hypothesis (53). Therefore, this CD39 finding has translational potential regardless of the causal direction of this interaction. We performed preliminary experiments using POM-1 to metabolically inhibit CD39. While we found no differences in bacterial burden between treatment and control groups, we did find POM-1 influenced cytokine expression during *Mtb* infection.

The functional state of T cells present in the lung may play a key role in the outcome of *Mtb* infection. However, it is not precisely clear how their functional state influences the control of *Mtb* growth. PD-1 is another inhibitory receptor that has been associated with *Mtb* bacterial load and *Mtb* specific CD4 T cells, however, PD-1 blockade treatment increases the incidence of TB reactivation (54). Moreover, mice deficient in PD-1 are extremely sensitive to *Mtb* infection and quickly succumb to death (55). We did not find the high CD39 expressing populations of T cells to also have high PD-1 expression.

While we found POM-1 treatment did not affect the bacterial burden in *Mtb* infected mice, this does importantly suggest that inhibition of CD39 does not increase the sensitivity of mice to *Mtb* infection. While the exact mechanism of action and knowledge of how immune cells will respond to inhibiting CD39 is still limited, one possible mechanism is that increased extracellular ATP levels from inhibiting ATP degradation could result in inflammasome-mediated release of pro-inflammatory cytokines including IL1β, which supports effector T cell and natural killer cell-mediated cytotoxicity (56). We found that POM-1 treated mice did have a trend of sustained increased Th1 cytokine expression in combination with an increase in IL1β expression up to 4 weeks post treatment. Taken together, our preliminary results showing that POM-1 affects cytokine expression albeit not bacterial burden in the *Mtb* infected lung indicate that CD39 may be a biomarker of disease severity rather than a contributor to disease severity. However, considerable work is required before we can conclude this finding. Moreover, our data supports further investigation into the inhibition of CD39 as a combination therapy with anti-TB antibiotics for the potential to reduce the length of treatment required to clear the bacteria.

In summary, our CyTOF investigation obtained a more detailed and expansive analysis of cellular composition in the mouse model that reflects relevant findings to TB disease in humans. Our analysis also defined functional cell states that could be a promising new target for treatment of TB. Our work highlights the complexity and dynamism of the cellular environment during various *Mtb* infection outcomes and exemplifies the importance of developing new models to interrogate the factors driving *Mtb* persistence or control.

Our study describes a mouse model to identify immune mechanisms that correspond to *Mtb* infection severity. The lung environment is highly dynamic during a *Mtb* infection and identification of relevant *Mtb*-specific immune events could be time and location dependent. The immune landscape analyzed here is from one timepoint after stabilization of infection and induction of the adaptive immune response and it is likely early immune events during the initial establishment of infection were missed. Further profiling of additional timepoints post infection along with analysis of vaccinated versus unvaccinated mice will provide greater insight into induced protective immunity and biomarkers of disease progression.

Our CyTOF experiments investigated immune cells at single-cell resolution, however our experiments do not spatially resolve lung immune cells. C57BL/6 mice do not form caseous granulomas in the lungs, which are typically found in human TB disease, therefore it may be difficult to translate spatial immune cell data from our experiment to human TB disease (11). CyTOF analysis focuses on the relative proportion of cells within each sample based on protein expression. For example, if we see a large infiltration of T-cells in the *Mtb* condition, we may see a small decrease in the other cell-types as the total proportion in each sample must sum to 100%. To account for the lack of spatial resolution, focus on protein data, and cell-type-proportion focused compositional analysis, we performed RNAscope analysis for mRNA associated with select proteins to improve spatial resolution and investigate whether we see findings at the protein and mRNA level. Indeed, we observed similar trends for select cell-type proportions between all three groups, supporting our conclusions.

CyTOF is also limited in that immune panels are pre-selected to investigate certain cell-types and cell-states in analyses. The advantages of CyTOF are the strong degree of multiplexing, the focus on protein expression, and that the number of cells collected per sample are an order of magnitude greater than single-cell transcriptomics. Future work applying single-cell transcriptomics to the immune subtypes we have identified in this study will allow us to find genes and pathways driving the dynamic cell-states that we observed.

We performed the first comprehensive, immune-system profiling of different TB disease states in the mouse lung. Our study reflects many previously discovered immune programs in response to *Mtb* infection within a single experimental context while indicating CD39 as an important biomarker for disease severity. Simultaneously, we characterized the mutant strain Δ*lsr2*, which shows similar attenuation to BCG while inducing a lung immune system profile reflecting an intermediate between *Mtb* and BCG. Our work will contribute to understanding of TB disease while providing candidates for TB biomarkers and a candidate for a live attenuated vaccine strain.

## Acknowledgements

We wish to thank the excellent staff at the Division of Comparative Medicine and the Toronto High Containment Facility. We also wish to acknowledge Ming Li for his help with the animal work. The study was supported by Canadian Institutes for Health Research Funds PJT-156261 and PJT-186285 (J.L.). Animal figures were created with BioRender.com.

## Author contributions

Conceptualization J.W. and J.L.; Methodology J.W., D.S., W.X., B.G. and K.B.; Investigation J.W., D.S., Y.Q., J.G., W.X., K.B. and B.G.; Visualization J.W. and D.S.; Funding acquisition J.L.; Project administration J.W.; Supervision D.B. and J.L.; Writing – original draft J.W., D.S. and J.L.; Writing – review and editing J.W. and J.L.

## Methods

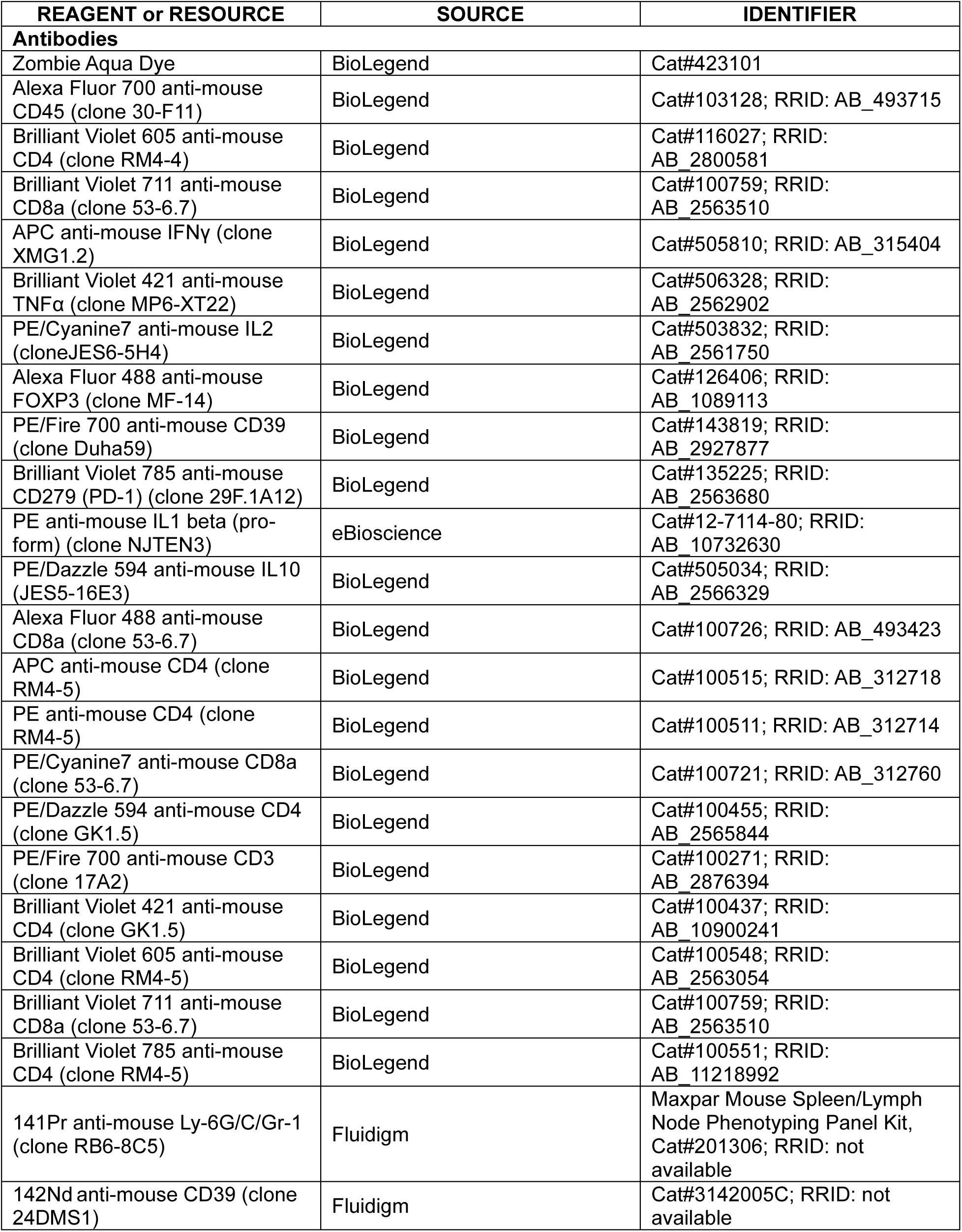
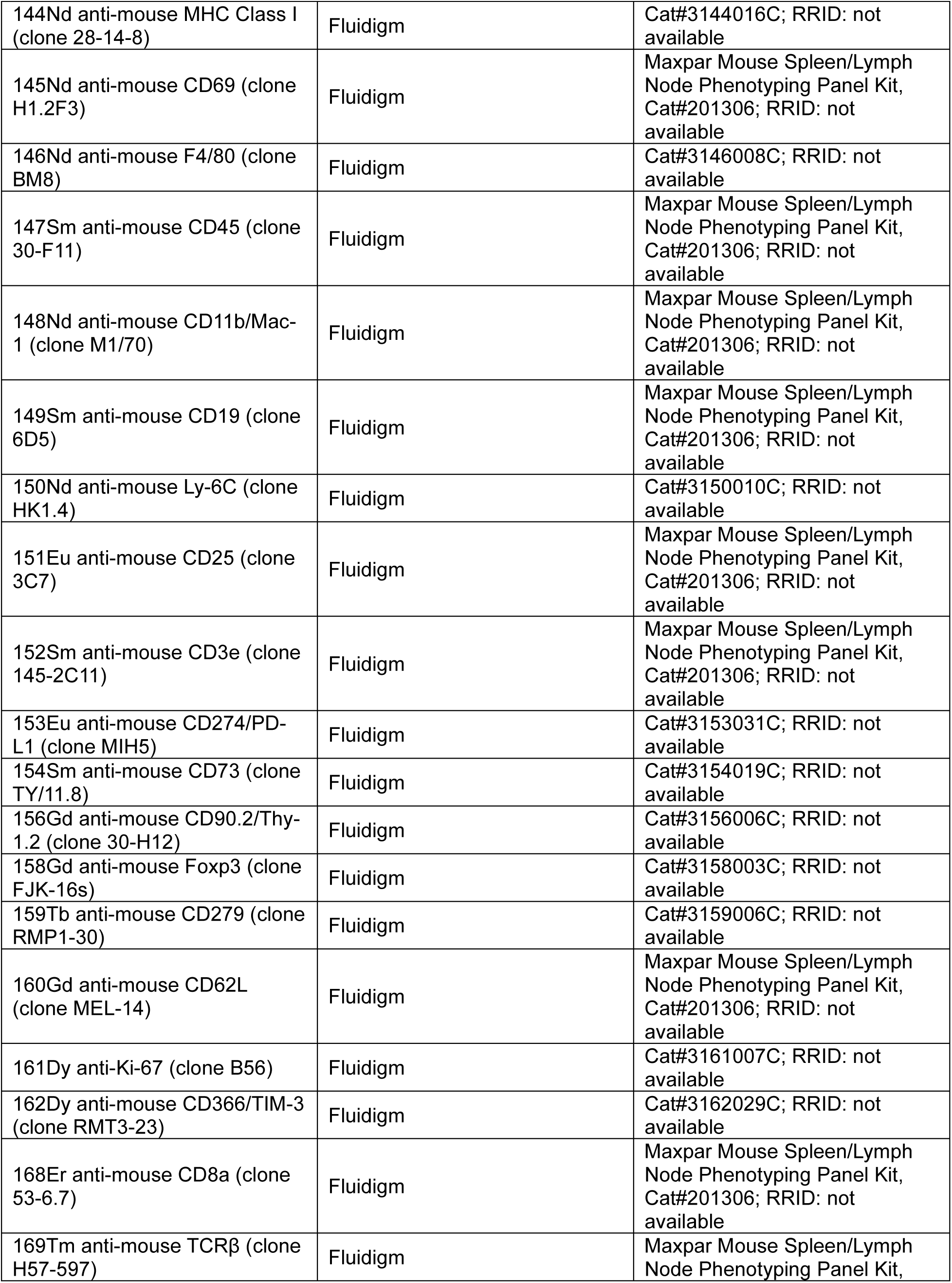
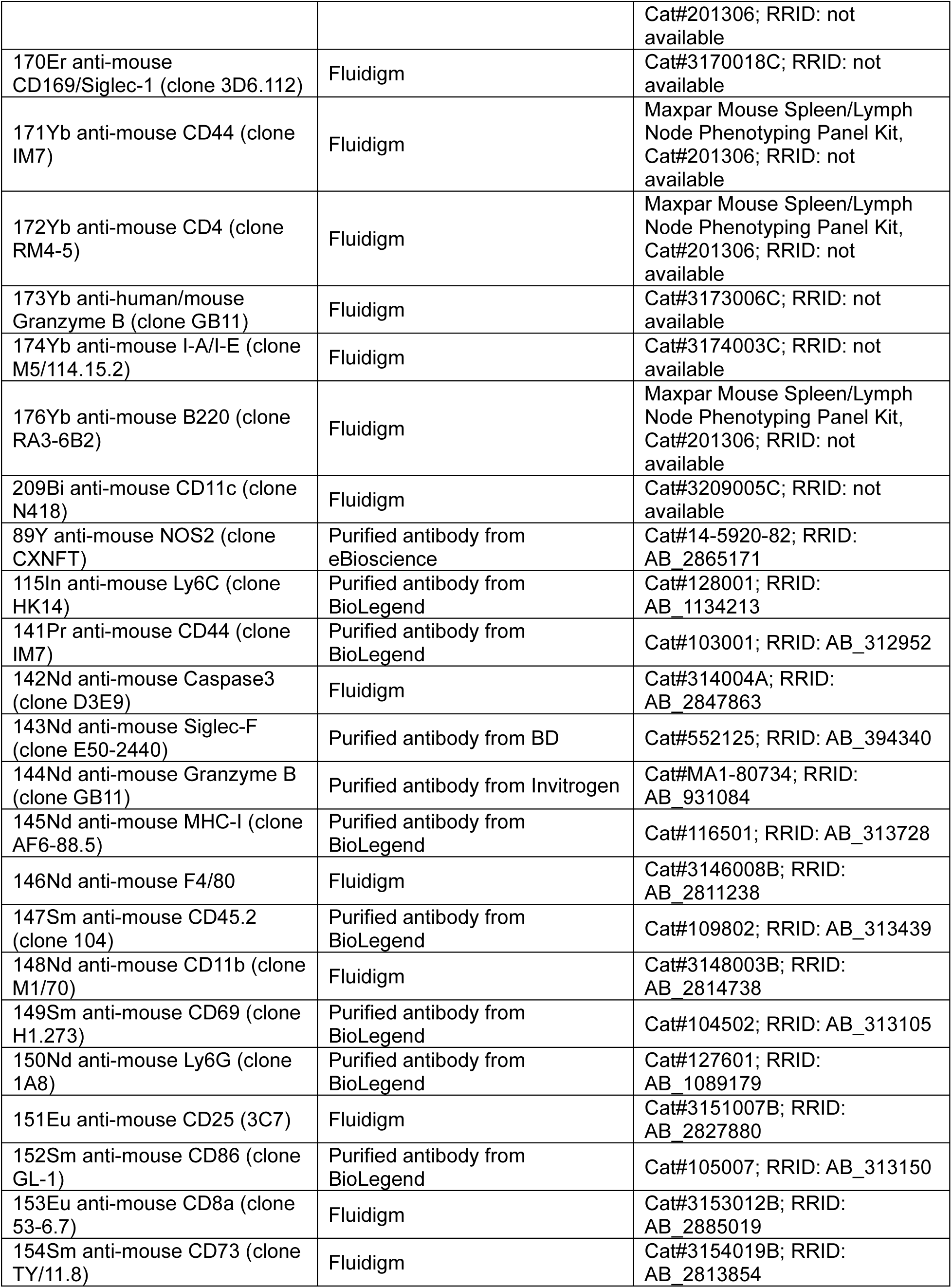
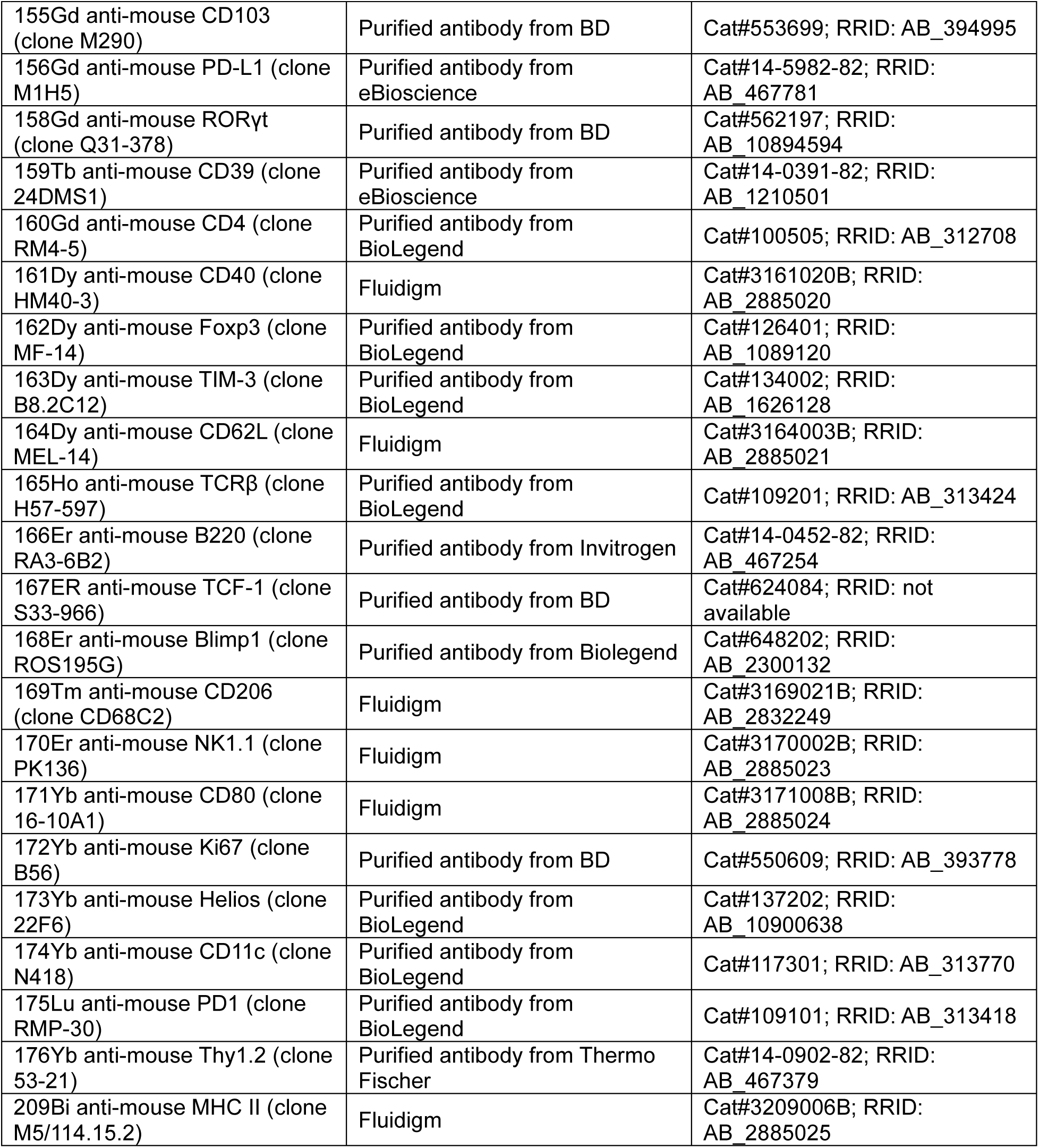

### Animals

C57BL/6 mice between 6-10 weeks of age were purchased from Charles River Laboratories. Female outbred Hartley Guinea pigs (200-250g) were purchased form Charles River Laboratories. Experiments involving *Mycobacterium tuberculosis* were conducted in the high containment facility, biosafety level III laboratory at the University of Toronto. Animal studies and procedures were approved by the University of Toronto Animal Care Committee and performed in accordance with the committees’ ethical standards.

### Bacterial strains and culture conditions

*Mycobacterium tuberculosis* H37Rv, *Mtb* Δ*lsr2*, and *M. bovis* BCG-Japan were prepared and titrated as previously described (57). *Mtb* strains were grown at 37°C in Middlebrook 7H9 broth (BD) supplemented with 0.2% glycerol, 10% albumin-dextrose-catalase, and 0.05% Tween 80 or on 7H10 agar supplemented with 1% casamino acids (Life), 0.5% glycerol, and 10% albumin-dextrose-catalase. BCG was grown at 37°C in Middlebrook 7H9 broth (BD) supplemented with 0.2% glycerol, 10% albumin-dextrose-catalase, and 0.05% Tween 80 or on 7H10 agar supplemented with 1% casamino acids (Life), 0.5% glycerol, and 10% oleic acid-albumin-dextrose-catalase. Hygromycin (BioShop) was added at a concentration of 75ug/mL for *Mtb* Δ*lsr2*.

A *lsr2* deletion mutant of *Mtb* was generated using a previously described TM4 phage-mediated specialized transduction, where the *lsr2* gene in *Mtb* H37Rv was replaced with a hygromycin-resistance cassette (25). Briefly, a targeted disruption of *lsr2* (Rv3597c) in *Mtb* H37Rv was made using an allelic exchange substrate by PCR amplifying the up and downstream DNA flanking the *lsr2* gene. The primer pairs used to PCR amplify the upstream and downstream flanking fragments for the allelic exchange substrate were (5’-CGGCTTCCATAAATTGGGCAGCTGGATCACCTGCTGGCGCAC-3’ and 5’-CGGCTTCCATTTCTTGGCATTTGGCTACCGGCGCCCAGGCGA-3’) and (5’- CGGCTTCCATAGATTGGTGGCTTACCCTCGCGTTTCTTCCTGTG-3’ and 5’-CGGCTTCCATCTTTTGGGGTGAAGAGATCACACCGCAGACG ACG-3’). The resulting PCR products were digested with the restriction enzyme PfIMI and ligated with the 1600 bp and 1760 bp fragments of the p0004 plasmid pretreated with the same enzyme to generate pKO-lsr2. pKO-lsr2 and phLR phasmid DNA were digested with PacI, ligated and packaged using the MaxPlax Lambda packaging extract (Epicentre), followed by transduction into *E. coli* NM759. The resulting phLR/pKO-lsr2 phasmid DNA from the transductants was electroporated into *M. smegmatis* mc2155 and plated for mycobacteriophage plaques at the permissive temperature of 30 °C. A high-titre phage lysate, prepared from one temperature-sensitive phage plaque, was used to infect H37Rv at the non-permissive temperature of 37°C as described previously (25). Hygromycin-resistant colonies were selected at 6-weeks post-transduction and confirmed by Southern hybridization.

### C57BL/6 mice infection with mycobacterium

For aerosol infections, 100-500 colony-forming units (CFUs) of *Mtb* H37Rv, *Mtb* Δ*lsr2*, or *M. bovis* BCG-Japan, per mouse was delivered using a Glas-Col nebulizer Inhalation Exposure System (57). Delivery dosage was confirmed by quantifying CFU in the lungs 1 day after exposure in control mice. At various time points after infection (2, 3, 4, 5, 6, 8 or 12 weeks) lungs and spleen CFUs were determined. Lungs and spleen were homogenized in PBS and tissue homogenate was serial diluted and coated on 7H10 agar plates. CFUs were counted on the plates after a 4-week incubation at 37°C. Tissues were also collected for flow cytometric and histological analysis.

### POM-1 treatment

Mice were aerosol infected with *Mtb* as described above. 3 weeks post infection, mice were intraperitoneally administered 5mg/kg POM-1 diluted in sterile PBS daily for 14 days. Control mice were administered sterile PBS only.

### Tissue isolation

Lung and spleen tissue was isolated as previously described (24). Lungs and spleen were minced into fine pieces then digested in RPMI 1640 medium without calcium/magnesium (Gibco) containing 10% FBS, 1% HEPES (Life), 1 mg/mL Collagenase from Clostridium histolyticum (Sigma) and 0.15 mg/mL DNase I (Sigma) on a shaker at 37°C for 1 hour with the lungs or 30 min with the spleen. Digested tissue was filtered through a 70 μm cell strainer to obtain single-cell suspensions and an aliquot was plated on 7H10 agar plates. CFUs were counted after a 4-week incubation at 37°C. The remaining lung and spleen single cell suspensions were red cell-lysed and then processed for flow cytometry.

To isolate leukocytes, after obtaining the single-cell suspensions samples were centrifuged with a 40/80% Percol (GE Healthcare Life Sciences) gradient and leukocytes were collected from the interface.

### Time-of-Flight mass cytometry (CyTOF)

Details for each antibody used for CyTOF are listed in Table 2.1. Directly conjugated antibodies were purchased from Standard Biotools. Single cell suspensions from individual samples were washed with PBS and stained with 1μM Cisplatin in PBS for 5min prior to quenching with Maxpar® Cell Staining Buffer and FBS underlay. Samples were then barcoded according to manufacturer’s instructions (Standard Biotools Cell-ID™ 20-Plex Pd Barcoding Kit) prior to being combined. Combined samples were resuspended in Maxpar® Cell Staining Buffer containing metal-tagged surface antibodies and Fc block (CD16/32; Bioxcell) for 30 minutes at 4°C. Cells were then washed, permeabilized and stained with metal tagged intracellular antibodies for 30 minutes at room temperature. Fixing and permeabilizing cells was done using the eBioscience Intracellular cytokine buffer kit and the eBioscience Foxp3 staining kit. Finally, cells were incubated overnight in PBS containing 0.3% (w/v) saponin, 1.6% (v/v) paraformaldehyde (Polysciences Inc) and 50nM Iridium (Standard Biotools). Cells were analyzed on a Helios2 mass cytometer (Standard Biotools) at The Hospital for Sick Children Center for Advanced Single Cell Analysis.

### Flow cytometry and intracellular cytokine re-stimulation

Details for each antibody used for flow cytometry are listed in Table 2.1. Cells were stimulated with phorbol 12-myristate 13-acetate and ionomycin and incubated with protein transport inhibitors Brefeldin A and Monensin for 5 h. Fixing and permeabilizing cells was done using the eBioscience Intracellular cytokine buffer kit and the Foxp3 staining kit (eBioscience). All antibody dilutions and cell staining were done using PBS containing 1% fetal calf serum and 2.5 mM EDTA. Zombie Aqua due was used to exclude dead cells from analyses.

Samples were analyzed on FACSSymphony^TM^ A5 (BD Bioscience) at the University of Toronto, Temerty Faculty of Medicine Flow Cytometry Core facility. Data was analyzed using Flow Jo Software v10.10 (BD FLowJo).

### Histological and RNA in situ hybridization analysis

A portion of the left lung and spleen was fixed for a minimum of one month in 10% neutral buffered formalin (Sigma-Aldrich) and paraffin embedded. The paraffin blocks were sectioned into 4 μm thick slices and were stained with hematoxylin and eosin (H&E) for histological analysis. The H&E staining was performed at the Centre for Phenogenomics in Toronto.

RNA in situ hybridization was performed using the RNAscope Multiplex Fluorescent v2 Assay (Advanced Cell Diagnostic), according to manufacturer’s instructions. Briefly, the paraffin blocks were continuously sectioned into 4 μm thick slices and mounted onto slides (Fisherbrand). The slides were baked at 60°C for 1 hour and then deparaffinized using xylene and ethanol. The samples were then treated with hydrogen peroxide for 10 min at room temperature. Target retrieval was done using RNAscope Target Retrieval Solution and then the samples were incubated with Protease Plus solution for 30 min at 40°C. Slides were incubated with the target probes cd3e (Advanced Cell Diagnostic, 314721), entpd1 (Advanced Cell Diagnostic, 475761), and IS6110 (Advanced Cell Diagnostic, 1208531) for 2 h at 40°C.

After hybridizing with the probes, signals were detected using TSA Vivid Fluorophore 520, TSA Vivid Fluorophore 570, and TSA Vivid Fluorophore 650. The slides were mounted with VectaMount Permanent Mounting Medium (Vector Laboratories). Whole-slide high resolution fluorescent scans were performed at 20x magnification with 3DHistech Slide Scanner at the Imaging Facility at the Hospital for Sick Children. DAPI, Cy3, Cy5, and GFP channels were used to acquire images. Images were processed and automated image analysis was performed with HALO v4 software (Indica Labs) at The Advanced Optical Microscopy Facility, University Health Network. Quantitative image analysis of RNAscope whole-tissue images was performed to determine the total number of *entpd1*, c*d3e*+, and IS6110+ cells using the classifier and FISH module v3.2.3 within HALO software.

### CyTOF data processing

In this study, two CyTOF datasets were generated, the following workflow was applied to each dataset individually. First, the output fcs files were debarcoded using the “CATALYST” R package (58,59). Specifically, the flow cytometry standard (.fcs) file generated by each CyTOF experiment was loaded into R using the read.FCS() function in the “flowCore” R package (60). The “debarcoding_gene_template.csv” was loaded, which represents a sample-by-mass matrix that allows us to extract the unique barcode assigned to each cell during the CyTOF experiment. The prepData() function in CATALYST with default parameters was used to filter non-mass channels, perform an arcsinh-transformation (cofactor = 5) (58,59), and generate a SingeCellExperiment object, an S4 object built to be compatible with many single-cell ‘omics R packages (61). Then, assignPrelim() was used to assign each cell to its sample of origin before filtering cells that were not assigned a unique barcode (58,59). Then the estCutoffs(), plotYields(), and applyCutoffs() functions were used to estimate barcode distributions, plot distributions, and filter cells whose distributions suggest technical artifacts such as doublets, respectively (58,59). Normalized and debarcoded sample files were saved as their own .fcs files.

For each dataset, data normalization, visualization, and quality control was performed manually using Flow Jo Software v10.10 (BD FLowJo) and by adapting the “cytofWorkflow” R package and module to our experimental design (62). This workflow predominantly relies on the “flowCore”, “CATALYST”, and “diffCyt” R packages (58,59,63). The following analysis was applied to each dataset individually. Debarcoded .fcs files for each sample were loaded into R using the “read.FCS” function before being converted into a flowSet object, which was designed by the “flowCore” R package to work with many .fcs files simultaneously (60). Then, the mass channel and mass key between each antibody to the biological symbol of the antibody was assigned (e.g., Tag 142Nd = CD39), combined the clinical metadata for each debarcoded sample, and generated a SingleCellExperiment object with the prepData() function (60). Quality control for these combined and normalized data was performed with the plotExprs() function, plotCounts(), and pbMDS() functions, which plot normalized protein intensity for each sample, plot the number of cells collected for each sample, and plot sample distribution based on cell-type specific protein expression profiles, respectively (60). Lastly, the plotNRS() function was used to visualize non-redundancy scores, which measure protein variance between cell types and conditions (60).

Normalized and quality-controlled data were then clustered to generate cell types. All proteins were used to cluster cells, rather than exclusively proteins profiled to capture cell types. All proteins were included in data clustering because including cell-state proteins allowed for empirically generated cellular subclusters that are driven by cellular activation, exhaustion, or apoptosis (e.g., a cluster defined by CD44 and CD4 to represent activated CD4 T cells) (58,59). Cells were clustered in each dataset using the cluster() function in the “CATALYST” R package, which first generates a 10 by 10 grid of clusters using FlowSOM (64), defined as self-organized map clustering optimized for CyTOF data. The cluster() function then groups these 100 clusters into 30 metaclusters using the ConsensusClusterPlus() R package (65), which combines K-means and hierarchical clustering. A heatmap that plots protein expression to each cluster using the “plotExprHeatmap” was then generate and to manually assign cell-types to each cluster (e.g., CD8a+, TCRβ+, CD4- = CD8 T cells). The relationship between protein expression, cluster ID, and sample-level cell-type proportions was visualized with the plotMultiHeatmap”. Lastly, dimed UMAP (66) dimensionality reductions was performed with the runDR() functions. For computational efficiency, 1800 randomly selected cells from each sample were used to generate UMAPs. UMAPs showing cell-type, cluster type, and protein expression are all plotted with the plotDR() function. With clusters manually mapped to cell-types and clusters visualized, the cell-type ID to each cell in the SingleCellExpression object was added using the mergeClusters() function. In each dataset, a small population of cells was found that were characterized by non-specific protein binding. These cells were labelled as “Undefined” and removed from downstream analysis. Similarly, a small population of cells that were defined by contradicting cell-type proteins (e.g., CD4, B220) were labelled as “Doublets” and removed from downstream analysis.

With cells normalized, assigned to their sample of origin, and assigned a cell-type identity (e.g., 100 clusters assignment, 30 clusters assignment, cell-type assignment), cluster/cell-type proportions was calculated using the diffcyt() R package with the “DA” module (63). Pseudobulked cluster/cell-type specific gene expression was also generated using the diffcyt() R package with the “DE” module (63). The final SingleCellExperiment, cell-type expression matrix and cell-type proportion matrix for each dataset were saved for dataset-specific downstream analysis.

### Differential protein and cell-type analysis from CyTOF data

A series of pairwise cell-type specific differential protein expression and differential cell-type abundance analyses was performed to calculate cell-type specific immune responses to each treatment. Cell-type specific differential abundance between each group (i.e., BCG vs. Δ*lsr2*, BCG vs. Δ*lsr2* vs. *Mtb*) was performed with the “diffcyt” package, using the “DS” analysis_type, the “diffcyt-DS-LMM” analysis method, and comparing protein abundance at the level of the cell-type (63). The “diffcyt-DS-LMM” uses a linear mixed model to compare protein abundance with parameters tuned for CyTOF data (63). Proteins whose median cell-type specific expression was less than 1 were excluded. Differentially active makers were selected with an FDR adjusted p-value <0.05. Cell-type protein abundances calculated from the “diffcyt” R package (63) were compared using a one-way ANOVA with three groups (i.e., BCG, *Mtb*, and Δ*lsr2*) and a Tukey’s HSD test. Cell-types whose median proportion was < 1% across each sample were excluded. Differentially abundant cell-types were selected with an FDR-adjusted p-value <0.05.

For each differential analysis, protein or cell-type differences with an FDR-adjusted p-value <0.05 were classified into a state. Differential proteins or cell-types between all three conditions were classified as “CFU”. Differential proteins or cell-types between BCG and each condition, but not between *Mtb* and Δ*lsr2* were classified as “MTB”, Differential proteins or cell-types between *Mtb* and each condition, but not between BCG and Δ*lsr2* were classified as “ATTN”. Differential proteins were organized by cell-type and condition and labelled by the state before being plotted into a heatmap with the Pheatmap R package (https://cran.r-project.org/web/packages/pheatmap/pheatmap.pdf). Differential cell-types were organized by condition, labelled by state, and plotted into a heatmap with the Pheatmap R package.

### Quantification and statistical analysis

Graphs and statistics were generated with GraphPad Prism v10 (GraphPad Software). Error bars represent the standard error of the mean (SEM). Student’s t-test, one-way and two-way ANOVA with Sidak’s or Tukey multiple comparison test was used to determine statistical significance where appropriate. Significance is indicated by: * = p <0.05, ** = p <0.01, *** = p <0.001, **** = p <0.0001. ns = not significant.

