## Supplemental Figures for "High Dimensional Immune Profiling Reveals CD39 as a Correlate of Tuberculosis Disease Severity"

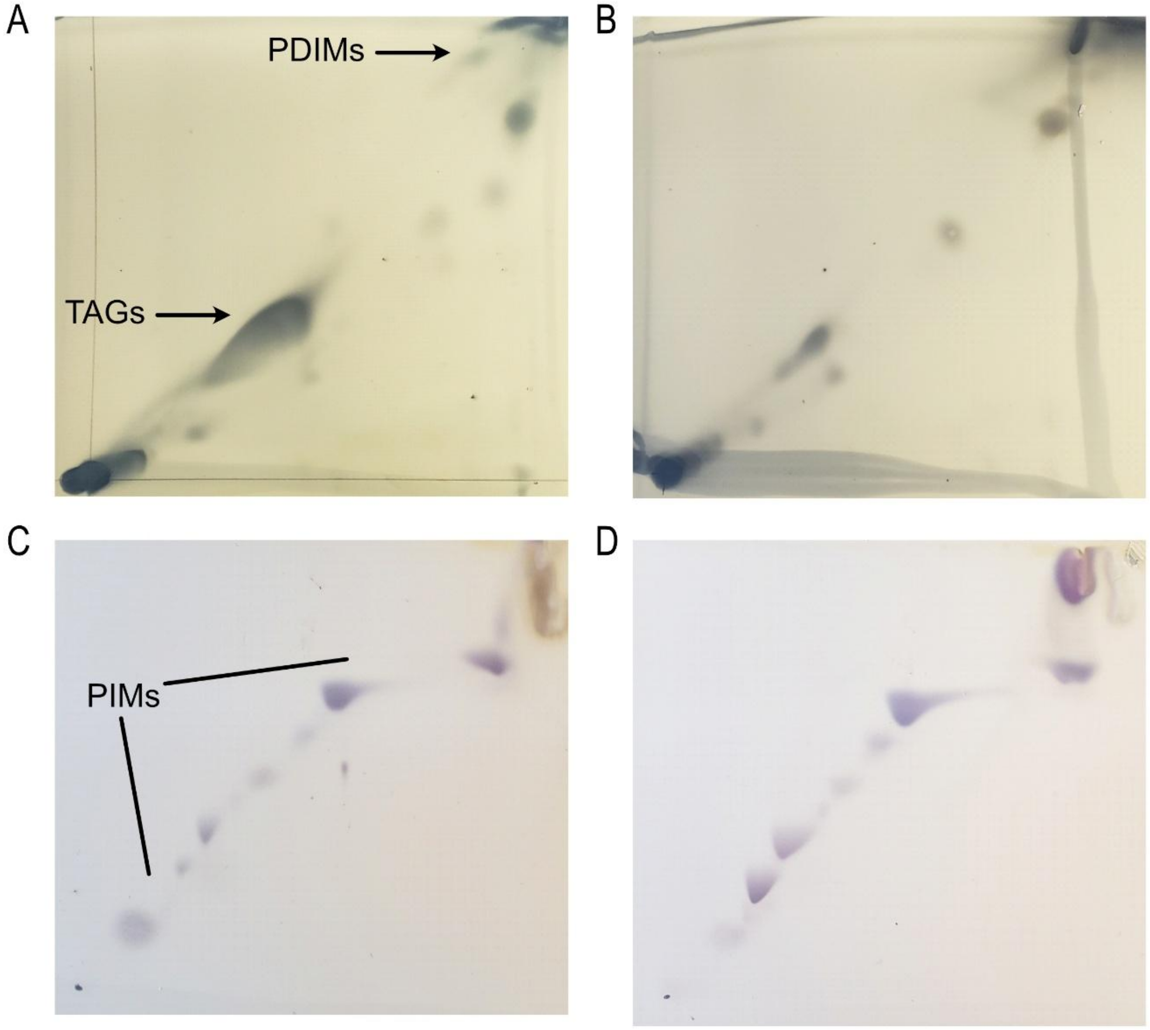


**Supplementary Figure S1 2D-TLC analysis of Δ*lsr2* cell wall lipids.** Apolar lipids of (A) *Mtb* H37Rv and (B) Δ*lsr2* were developed with the solvent system petroleum ether/ethyl acetate (49:1, v/v, 3x) in the first dimension and petroleum ether/acetone (49:1, v/v) in the second dimension, followed by charring with 5% phosphomolybdic acid. PDIMs: phthiocerol dimycocerosates. TAGs: triacylglycerol. Polar lipids of (C) *Mtb* H37Rv and (D) Δ*lsr2* were separated with the solvent system: chloroform/methanol/water (60:30:6, v/v) in the first dimension and chloroform/acetic acid/methanol/water (40:25:3:6, v/v) in the second dimension, followed by charring with α-naphthol. PIMs: phosphatidylinositol mannosides.

A

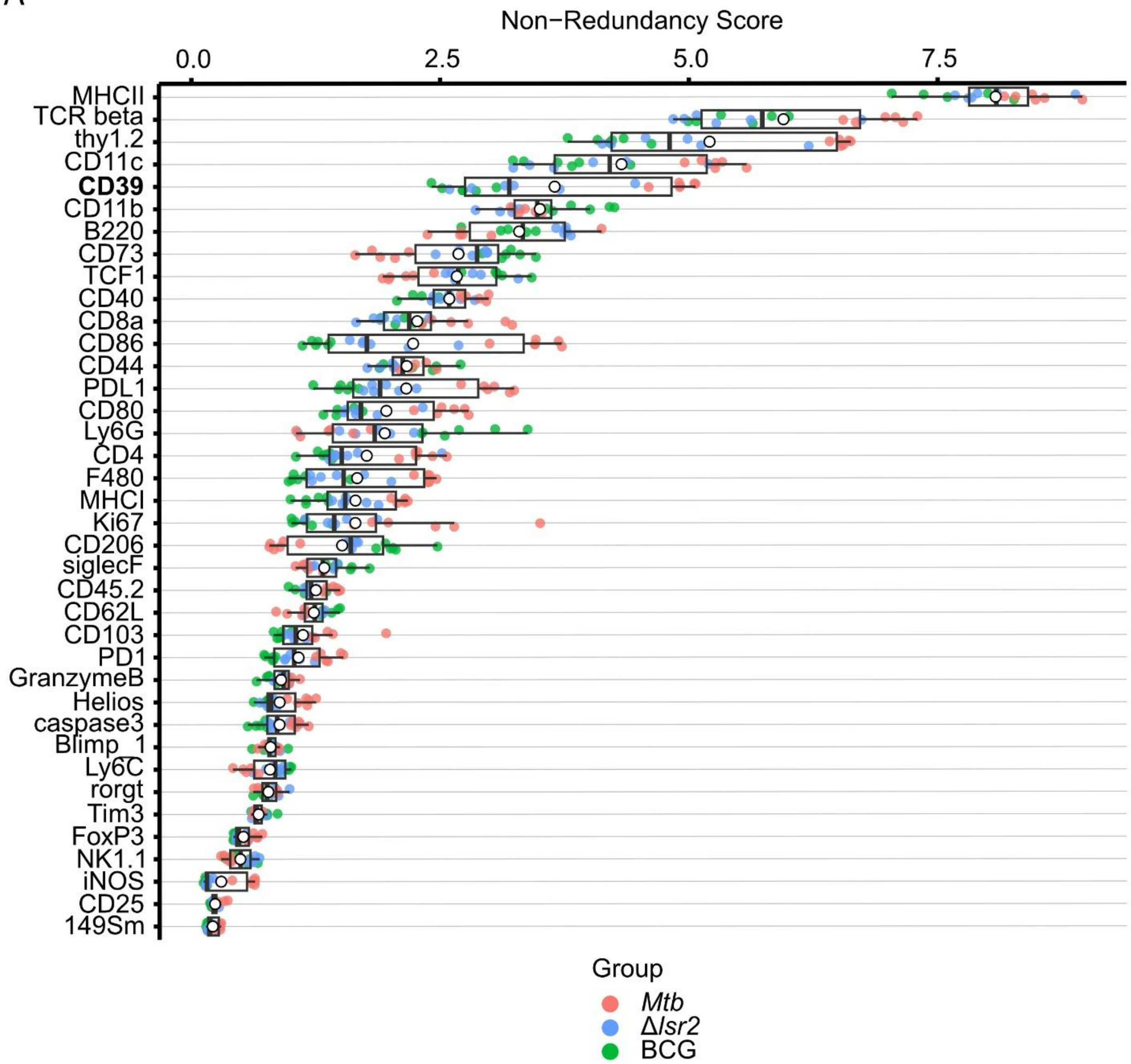


**Supplementary Figure S2 CD39 largely contributes to variation in the samples.** (A) Non-redundancy scores for all proteins and all samples in the dataset. The full-coloured points represent the per-sample non-redundancy scores, coloured by group. The empty black circles indicate the mean non-redundancy scores from all the samples. Proteins on the y-axis are sorted according to the decreasing average non-redundancy score

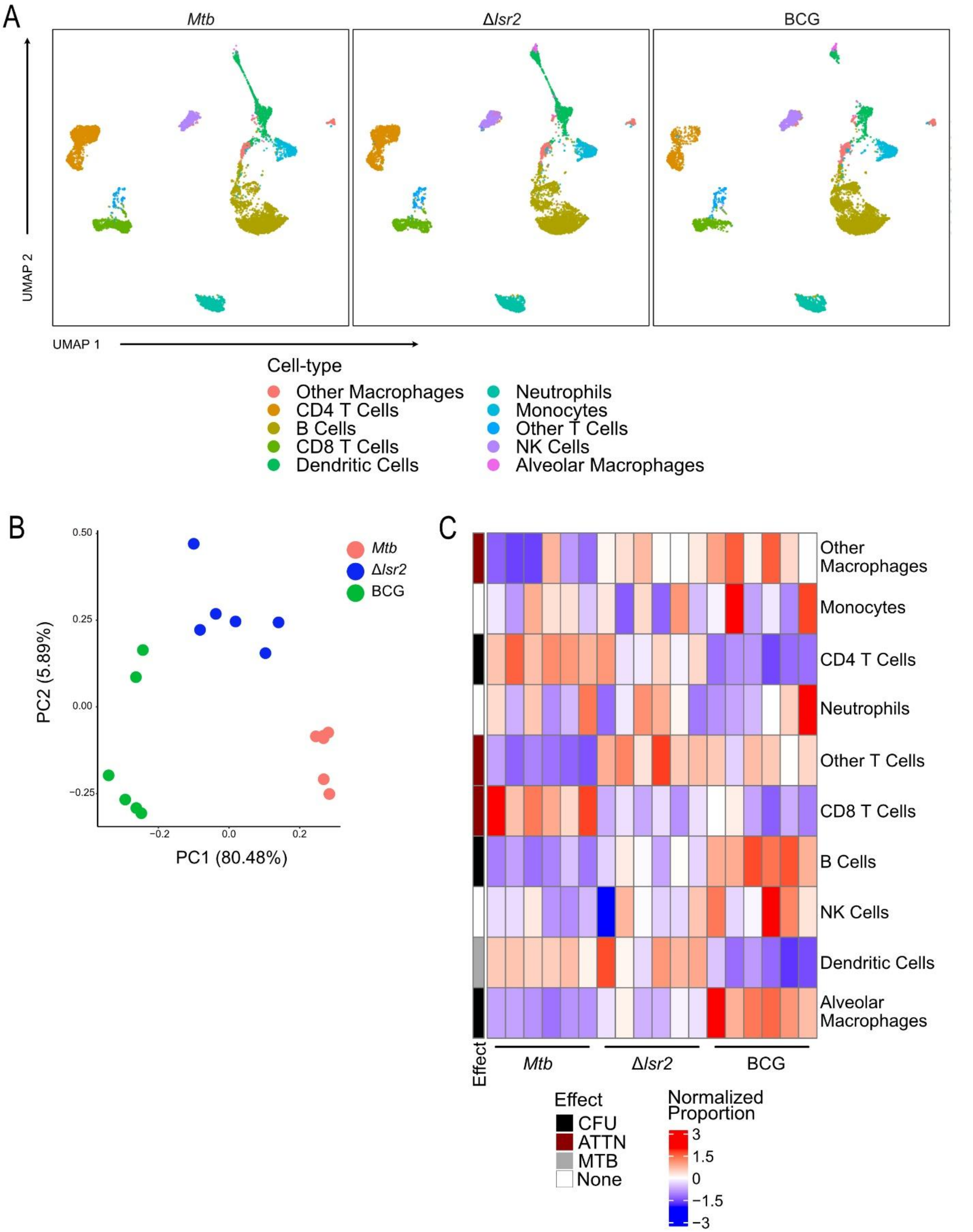


**Supplementary Figure S3 Lung-infiltrating cell analysis allows for better detection of the cell-type abundance and marker expression dynamics between lower bacterial burden groups, Δ*lsr2* and BCG.** 6 C57BL/6 mice per group were aerosol challenged with *Mtb* H37Rv, Δ*lsr2*, or BCG (~100 CFU/lung). Lung samples were perfused with PBS before being processed into single cell suspensions and analyzed with CyTOF. (A) Uniform manifold approximation mapping (UMAP) of cell-type specific protein expression. Panels are split by group. Cells are labelled by cell-type identified by manually annotating metaclusters based on their protein expression profiles. (B) Scatterplot visualizing the first two principal components of pseudobulked cell-type specific protein expression. Colours represent condition. (C) Heatmap represents differences in cell-type abundance between each treatment. Rows are different cell-types, columns are different samples, and heat represents row-normalized cell-type abundance. Rows are annotated by their state (i.e. CFU, ATTN, or MTB) for cell-type differences with an FDR-adjusted p-value <0.05. None = not significant.

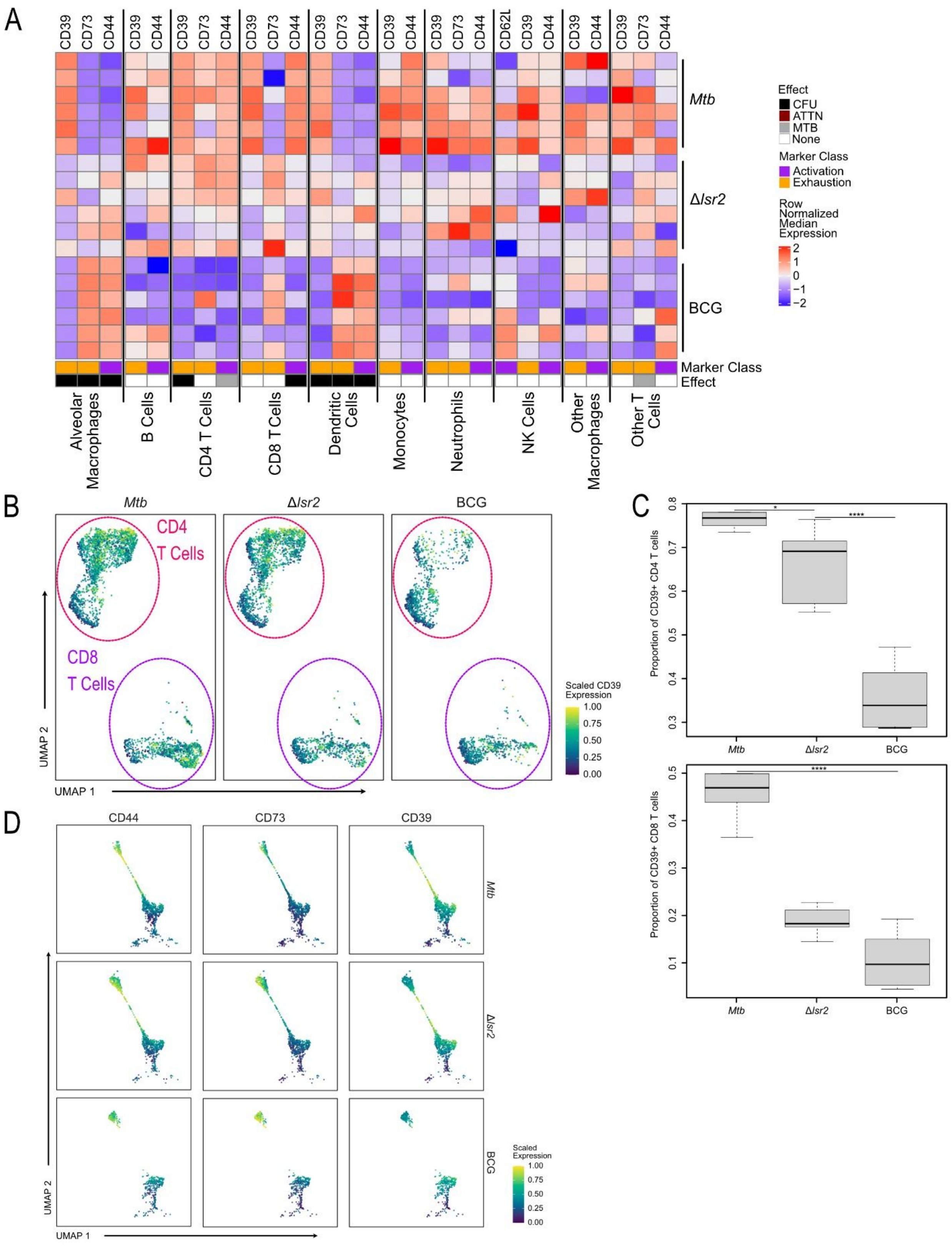


**Supplementary Figure S4 Lung-infiltrating cell analysis allows for better detection of marker expression dynamics between Δ*lsr2* and BCG.** (A) Heatmap represents differences in cell-state protein expression between each treatment in all immune cells. Columns represent cell-type proteins and rows represent samples. Cells are populated by row-normalized protein expression. Columns are annotated by cell-type, state (i.e. CFU, ATTN, or MTB), and protein class (i.e., activation, exhaustion etc.). State is shown for cell-type protein expression differences with an FDR-adjusted p-value <0.05. None = not significant. (B-C) A high proportion of CD4 and CD8 T cells express CD39 after an *Mtb* infection in the PBS-perfused lung. (B) UMAP of CD39 protein expression performed in CD4 and CD8 T cells. (C) Bar plots comparing the ratio of CD39+ CD4 T cells vs CD39- CD4 T cells and CD39+ CD8 T cells vs CD39- CD8 T cells within each group. Differences in cell-type CD39 positivity was measured using a one-way ANOVA and Tukey's HSD test, with each mouse as a biological replicate (*, p <0.05; **, p <0.01; ***, p <0.001; ****, p <0.0001; ns = not significant). (D) UMAP of differentially expressed proteins in dendritic cells in the PBS-perfused lung. Columns represent cell-type protein expression for CD44, CD73, and CD39. Rows are annotated by infection group.

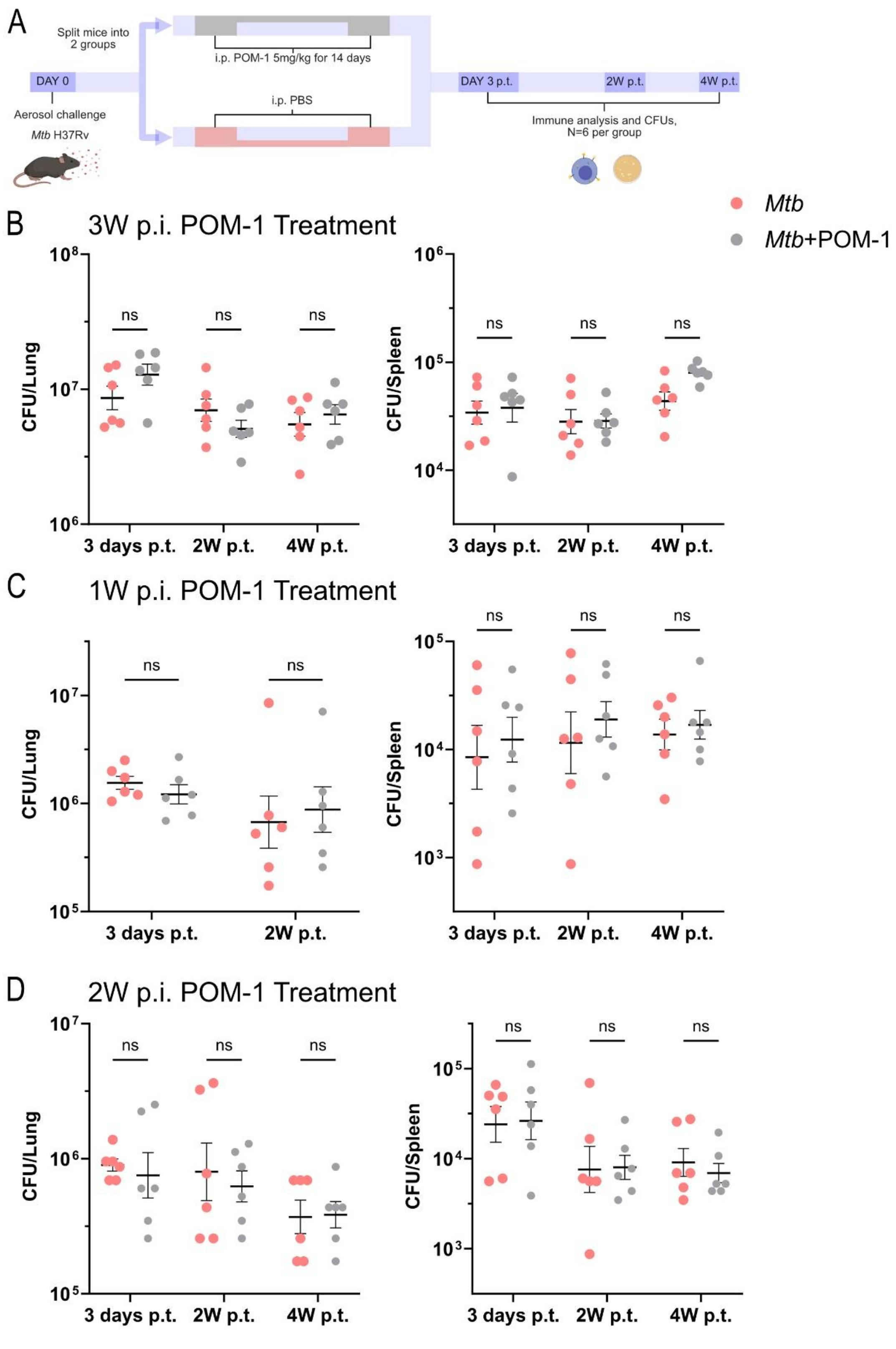


**Supplementary Figure S5 POM-1 treatment does not affect *Mtb* burden regardless of treatment initiation timing post infection.** (A) Schematic representation of experimental design. C57BL/6 mice were aerosol challenged with *Mtb* H37Rv (~100 CFU/lung) and after 1 week, 2 weeks, or 3 weeks were treated with either 5mg/kg of POM-1 or PBS as a control. *Mtb* burden in the lungs and spleen was determined at 3 days post treatment (p.t.), 2 weeks p.t. and 4 weeks p.t. (B) CFU data for POM-1 treatment initiated 3 weeks p.i. (C) CFU data for POM-1 treatment initiated 1 week p.i. (D) CFU data for POM-1 treatment initiated 2 weeks p.i. Data presented as mean ±SEM. Statistical significance was determined by two-way ANOVA with Sidak's multiple comparisons test (*, $p < 0.05$; **, $p < 0.01$; ***, $p < 0.001$; ****, $p < 0.0001$; ns = not significant).
